# Constitutive sodium permeability in a *C. elegans* two-pore domain potassium channel

**DOI:** 10.1101/2024.01.22.576648

**Authors:** O Andrini, I Ben Soussia, P Tardy, DS Walker, C Peña-Varas, D Ramírez, M Gendrel, M Mercier, S El Mouridi, A Leclercq-Blondel, W González, WR Schafer, M Jospin, T Boulin

**Affiliations:** University Claude Bernard Lyon 1, MeLiS, CNRS UMR 5284, INSERM U1314, 69008 Lyon, France; Neurobiology Division, MRC Laboratory of Molecular Biology, Cambridge, UK; Departamento de Farmacología, Facultad de Ciencias Biológicas, Universidad de Concepción, Chile; Center for Bioinformatics, Simulation and Modelling (CBSM) University of Talca, Chile; Department of Biology, KU Leuven, Leuven, Belgium

**Keywords:** Two-pore domain potassium channel, ion selectivity, molecular dynamics, electrophysiology, *C. elegans*

## Abstract

Two-pore domain potassium (K2P) channels play a central role in modulating cellular excitability and neuronal function. The unique structure of the selectivity filter in K2P and other potassium channels determines their ability to allow the selective passage of potassium ions across cell membranes. The nematode *C. elegans* has one of the largest K2P families, with 47 subunit-coding genes. This remarkable expansion has been accompanied by the evolution of atypical selectivity filter sequences that diverge from the canonical TxGYG motif. Whether and how this sequence variation may impact the function of K2P channels has not been investigated so far. Here we show that the UNC-58 K2P channel is constitutively permeable to sodium ions and that a cysteine residue in its selectivity filter is responsible for this atypical behavior. Indeed, by performing *in vivo* electrophysiological recordings and Ca^2+^ imaging experiments, we demonstrate that UNC-58 has a depolarizing effect in muscles and sensory neurons. Consistently, *unc-58* gain-of-function mutants are hypercontracted, unlike the relaxed phenotype observed in hyperactive mutants of many neuromuscular K2P channels. Finally, by combining molecular dynamics simulations with functional studies in *Xenopus laevis* oocytes, we show that the atypical cysteine residue plays a key role in the unconventional sodium permeability of UNC-58. As predicting the consequences of selectivity filter sequence variations *in silico* remains a major challenge, our study illustrates how functional experiments are essential to determine the contribution of such unusual potassium channels to the electrical profile of excitable cells.

**SIGNIFICANCE:** Potassium channels play a central role in modulating cellular excitability, particularly of neuronal cells. Their unique structure determines their ability to let ions pass selectively through cell membranes. The impact of pathological or evolutionary variations in this selectivity filter remains difficult to predict. Here, we reveal that UNC-58, a member of the two-pore domain potassium channel family of *C. elegans*, exhibits an unusual sodium permeability due to a unique cysteine residue in its selectivity filter. Our findings underscore the importance of functional studies to determine how sequence variation in potassium channel selectivity filters can shape the electrical profiles of excitable cells.

## INTRODUCTION

Potassium channels are an essential class of transmembrane proteins that allow the selective passage of potassium cations (K^+^) across biological membranes. Under physiological conditions, the opening of these channels allows the outward flow of K^+^ that drives membrane repolarization and participates in the maintenance of a hyperpolarized resting membrane potential. The high selectivity of the channels for K^+^ over sodium (Na^+^) is therefore a crucial feature that determines their role in shaping cellular excitability.

Comparison of K^+^ channel amino acid sequences, structure-function studies and the determination of the atomic structure of the bacterial KcsA channel have elucidated the mechanisms through which potassium channels facilitate the rapid conduction of K^+^ ions while simultaneously preventing the passage of smaller cations, such as Na^+^ (Doyle et al., 1998; Heginbotham et al., 1994, 1992; Shealy et al., 2003). The pore region of all K^+^ channels is constituted by the juxtaposition of four short amino-acid segments, called P-loops, and four inner transmembrane helices (see for reviews (Kim and Nimigean, 2016; Kuang et al., 2015; Mironenko et al., 2021; Roux, 2017)). To cross the channel from the intracellular side, K^+^ enters a cavity through the bundle formed by the four inner transmembrane helices. K^+^ ions remain hydrated in this vestibule. Potassium ions then proceed through the final section of the pore, i.e., the selectivity filter (SF), which is so constricted that a K^+^ ion must shed its hydration layer to gain entry. The sequence of the selectivity filter is highly conserved and consists, in most cases, of a stretch of five residues (TxGYG). These residues create four identical and equally-spaced binding sites for K^+^, called S1, S2, S3 and S4 (from the extracellular to the intracellular side). The S1, S2, and S3 sites are formed by the carbonyl oxygens from the backbone of the three residues xGY, while the S4 site is defined by the carbonyl of the threonine residue and the hydroxyl oxygen of its lateral chain. Each of these sites can stabilize a fully dehydrated K^+^ ion that is coordinated by four lower and four upper oxygen atoms. It is generally believed that K^+^ ions occupy either the S1 and S3, or the S2 and S4 sites, with water molecules filling the remaining positions. This arrangement averts repulsive electrostatic interactions between K^+^ ions. In the presence of an electrochemical driving force, water-K^+^ pairs move together through the selectivity filter. In contrast, the free energy barriers are higher for Na^+^ than K^+^ in the selectivity filter, which hinders sodium from efficiently passing through the pore (Kim and Nimigean, 2016; Kuang et al., 2015; Mironenko et al., 2021; Roux, 2017).

In contrast to the tetrameric structure of voltage-gated K_v_ and inwardly-rectifying K_ir_ potassium channels, two-pore domain potassium (K2P) channels are dimers of subunits, each featuring two distinct selectivity filter sequences (SF1 and SF2) (Figure 1A, B) (for review see (Natale et al., 2021)). SF1 and SF2 often differ in their amino acid composition. For instance, the tyrosine of the canonical TxGYG motif is generally replaced by a leucine or a phenylalanine in SF2. In addition, the linker between SF2 and the fourth transmembrane helix is often longer than the linker between SF1 and the second transmembrane helix (or linkers of other K^+^ channels). One might expect that these sequence variations would result in an asymmetric structure that would challenge the classical model of K^+^ selectivity filter organization. However, the elucidation of crystal structures for TWIK1 and TRAAK (Brohawn et al., 2012; Miller and Long, 2012) revealed a quasi-4-fold symmetry of the selectivity filter that preserves K^+^ coordination. Thus, the structures of K2P channel selectivity filters underscore the strong evolutionary pressures that ensure selective conduction of K^+^ through these channels.

**Figure 1.**
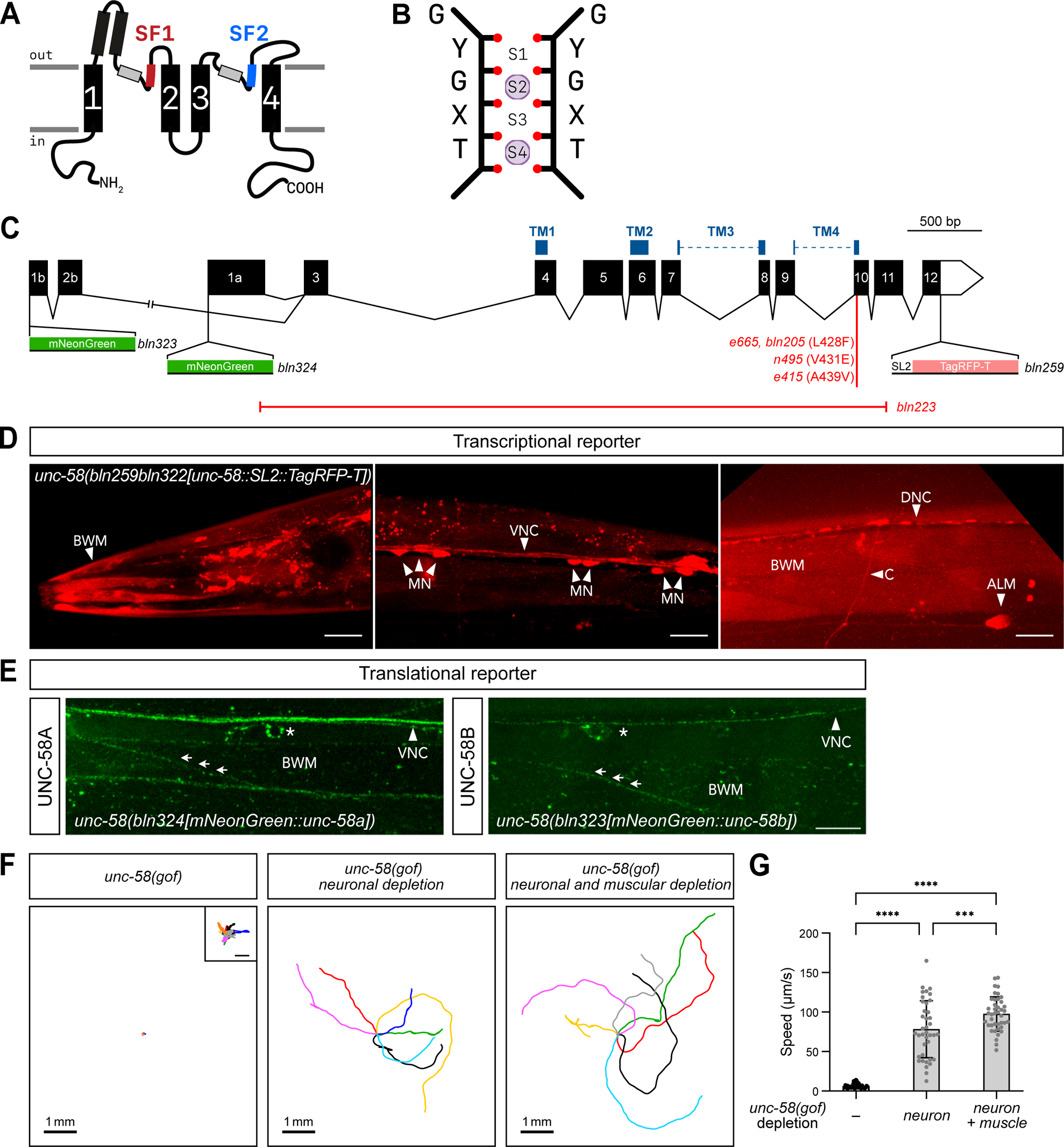
The two-pore domain potassium channel UNC-58 is expressed in neurons and muscle cells in *C. elegans*. **A** Schematic transmembrane topology of a two-pore domain (K2P) potassium channel subunit. Transmembrane segments are numbered “1” through “4”. The two selectivity filter regions (SF1 and SF2) precede the second and fourth transmembrane helix. A pore helix (light gray rectangles) precedes each selectivity filter. Functional channels are formed by the dimerization of two K2P subunits. **B** Schematic representation of a prototypical potassium channel selectivity filter. Four evenly-spaced ion binding sites (S1–S4) are formed by the carbonyl oxygens (red dots) of the TxGYG selectivity filter residues. Two ions are positioned in S2 and S4 (purple). **C** *unc-58* gene structure, position of transmembrane domains (TM1 through TM4, above exons), point mutations (*e665, bln205, n495, e415*) and deletion (*bln223,* red line), transcriptional (*bln259*) and translational N-terminal knock in alleles (*bln323, bln324*). Amino acid positions in reference to *unc-58* isoform b (transcript T06H11.1b.1). **D** *unc-58* transcriptional reporter. Left panel, anterior body wall muscles (BWM). Middle panel, ventral nerve cord (VNC) and cell bodies of ventral nerve cord motoneurons (MN). Right panel, ALM, commissures (C, arrowhead), and dorsal nerve cord (DNC). Scale bar, 10 µm. **E** *unc-58* translational reporter. Left panel, UNC-58A isoform, mNeonGreen was fused in frame upstream of exon 1a. Right panel, UNC-58B isoform, mNeonGreen was fused in frame upstream of exon 1b. Asterisk, neuronal cell bodies. Arrows indicated body wall muscle plasma membranes. Scale bar, 10 µm. **F** Displacement of worms over 50 seconds on nematode growth media. Depletion of UNC-58(gof) in neurons and muscle improves locomotion compared to depletion of UNC-58(gof) in neurons alone. Inset, tenfold-magnified view of *unc-58(gof)* tracks. Scale bar, 50 µm. **G** Average speed of *unc-58(gof)* (n = 49), neuronally- (n = 39), and neuronally- and muscle-depleted (n = 41) *unc-58(gof)* animals. Bar, average; whiskers, standard deviation. Each point represents the speed of an animal. One-way ANOVA, followed by Holm-Šídák’s multiple comparisons test, *** p<0.001, **** p<0.0001.

While K^+^ selectivity is generally a constant property of K^+^ channels, dynamic changes in K^+^ versus Na^+^ selectivity have been described for K2P channels (Chatelain et al., 2012; Chen et al., 2014; Liu et al., 2023; Ma et al., 2012, 2011). For example, in heterologous systems and in primary cultures of human cardiomyocytes, TWIK1’s selectivity switches from K^+^ to Na^+^ when the external concentration of potassium is decreased (Ma et al., 2011). This behavior has been linked to the presence of a threonine residue preceding the GYG in the SF1 sequence of TWIK1 (T<u>T</u>GYG). Mutation of this threonine to an isoleucine, which is found in most K2P channels at this position, restores K^+^ selectivity when external K^+^ concentration is lowered. Conversely, substituting the isoleucine found at the corresponding position in TASK3 with a threonine converts the purely K^+^ permeable channel into a channel permeable to Na^+^ when extracellular potassium levels are decreased. Interestingly, the same mutation does not alter the selectivity of the THIK1 K2P channel in low extracellular potassium, suggesting that this threonine is important for dynamic selectivity but not always sufficient (Ma et al., 2011). These findings highlight the difficulty of predicting selectivity changes in a given channel.

An interesting alternative to conventional structure/function approaches, which rely on point mutation engineering, is to capitalize on the inherent sequence diversity provided by evolution. The genome of the model nematode *Caenorhabditis elegans* contains 47 genes encoding K2P channel subunits, compared to 15 genes in humans. In several instances, nematode channels exhibit intriguing changes in their selectivity filter sequences.

The precise role of many nematode K2P channels has not been explored so far, but a few null and gain-of-function mutants have been characterized. Null mutations in nine K2P channels expressed in muscle or motor neurons lead to either no gross phenotype or to mild locomotor defects (de la Cruz et al., 2003; Lüersen et al., 2016; Zhou et al., 2022). In contrast, gain-of-function mutations often result in easily visible phenotypes. For instance, gain-of-function mutants of K2P channels expressed in skeletal muscles, such as SUP-9, TWK-18 or TWK-28 (Ben Soussia et al., 2019; de la Cruz et al., 2003; Kunkel et al., 2000; Peysson et al., 2023) cause muscle relaxation and a flaccid paralysis. Increased activity of K2P channels expressed in neurons of the motor circuitry, such as TWK-2, TWK-7 or TWK-40 (Yue et al., 2022; Zhou et al., 2022) similarly results in dramatic locomotor impairment and relaxed posture.

These locomotor defects stem from a decrease in neuromuscular excitability, a logical consequence of K^+^ channel hyperactivation. Remarkably, gain-of-function mutants of the UNC-58 K2P channel display a very different phenotype. These animals exhibit a rigid paralysis and a shortened body length, a phenotype consistent with hypercontracted body wall muscles (Ben Soussia et al., 2019; Kasap and Dwyer, 2021; Salkoff, 2006). Based on these observations, it has been proposed that the ion selectivity of UNC-58 might be altered (Kasap and Dwyer, 2021; Salkoff, 2006).

In this study, we have performed a detailed functional characterization of the *C. elegans* UNC-58 channel. We show that *unc-58* is expressed in muscle and neuronal cells. Using electrophysiology and *in vivo* calcium-imaging, we demonstrate that UNC-58 has an excitatory rather than inhibitory function. Consistently, molecular dynamics simulations and structure/function studies indicate that it constitutively conducts Na^+^ ions. Finally, we identify a residue in the SF1 selectivity filter sequence that is unique to UNC-58 and plays a critical role in the unusual permeability of this K2P channel.

## RESULTS

### The two-pore domain potassium channel UNC-58 is expressed in muscle and neurons

*unc-58* gain-of-function (*gof*) mutants exhibit a peculiar phenotype. They are hypercontracted, do not produce any forward or backward movement, but constantly rotate around their antero-posterior axis (Ben Soussia et al., 2019) (Supplementary movie SM1). Gain-of-function mutations identified in forward genetic screens harbor single amino acid changes in the fourth transmembrane domain. For example, the canonical *e665* mutation results in the substitution of a leucine at position 428 by a phenylalanine residue (L428F) (Figure 1C). In contrast, null mutants of *unc-58* are only moderately uncoordinated (Rawsthorne-Manning et al., 2022; Zhou et al., 2022).

These locomotor impairments suggested a role for the channel in the control of neuromuscular excitability. To determine the expression profile of *unc-58*, we generated a transcriptional reporter strain in which an SL2::TagRFP-T cassette was inserted at the 3’ end of the *unc-58* gene using CRISPR/*Cas9* genome editing (Figure 1C). Using this fluorescent reporter, we found that *unc-58* was broadly expressed in muscle cells and neurons (Figure 1D). We observed expression in striated body wall muscles and in almost all classes of ventral nerve cord motor neurons. We detected *unc-58* expression in all GABAergic D-type neurons, and in all cholinergic A and B-type motoneurons, except VA1 and DB1. We also detected weak expression in AS motoneurons and in the VC4 neuron (Figure S1 and Supplementary Table ST1). In addition, using well-characterized fluorescent neuronal markers, we could confirm *unc-58* expression in over 60 neurons using all types of neurotransmitters in the head and tail ganglia and belonging to different neuron classes (Figure S1 and Supplementary Table T1). Finally, we could detect *unc-58* in the GLR glial cells.

To visualize the subcellular localization of UNC-58, we engineered translational reporters by inserting the coding sequence of the fluorescent protein mNeonGreen into the first exon of two *unc-58* isoforms (Figure 1C). Both isoforms were observed in neurons and muscle cells, and were clearly visible at the membrane of muscle cells and in neurons of the nerve cords (Figure 1E).

To determine whether the locomotor defects of *unc-58(gof)* were due to its role in motor neurons and/or muscle cells, we performed tissue-specific degradation of UNC-58. We fused the ZF1 sequence (Armenti et al., 2014) to UNC-58, and expressed the ZIF-1 SOCS-box adaptor protein under the control of specific promoters for muscle (*myo-3^prom^*) or A and B-class motor neurons (*acr-2^prom^*) (Meng, 2022). Locomotion was clearly restored by depletion of UNC-58 L428F from neurons, and could be further improved by removal of UNC-58 from both muscles and neurons (Figure 1F, G and Supplementary movies SM2, SM3, and SM4). These results suggest that the *unc-58(gof)* phenotype has both muscle and motor neuron contributions.

### UNC-58 gain-of-function results in hyperactivity of ALM touch neurons and muscle cell depolarization

To assess the functional role of UNC-58 channels in neurons, we focused on the mechanosensory neuron ALM. ALML and ALMR are two of the six gentle touch receptor neurons of *C. elegans* (Chalfie et al., 1985). Well-controlled mechanical stimulation was applied with a glass probe, while we monitored calcium activity using a genetically-encoded fluorescent calcium indicator. To enable us to uncover increases in touch sensitivity, as well as decreases, we used a “short press”, a milder stimulation than the standard gentle touch protocol (Suzuki et al., 2003), such that wild-type worms responded only 62 % of the time. *unc-58* gain-of-function mutants showed increased response rates, responding 93 % of the time and higher average calcium responses (Figure 2A, B). Conversely, *unc-58* null mutants showed clearly reduced response rates (33%) (Figure 2B). Both results were inconsistent with the expected hyperpolarizing function of a potassium-selective channel.

**Figure 2.**
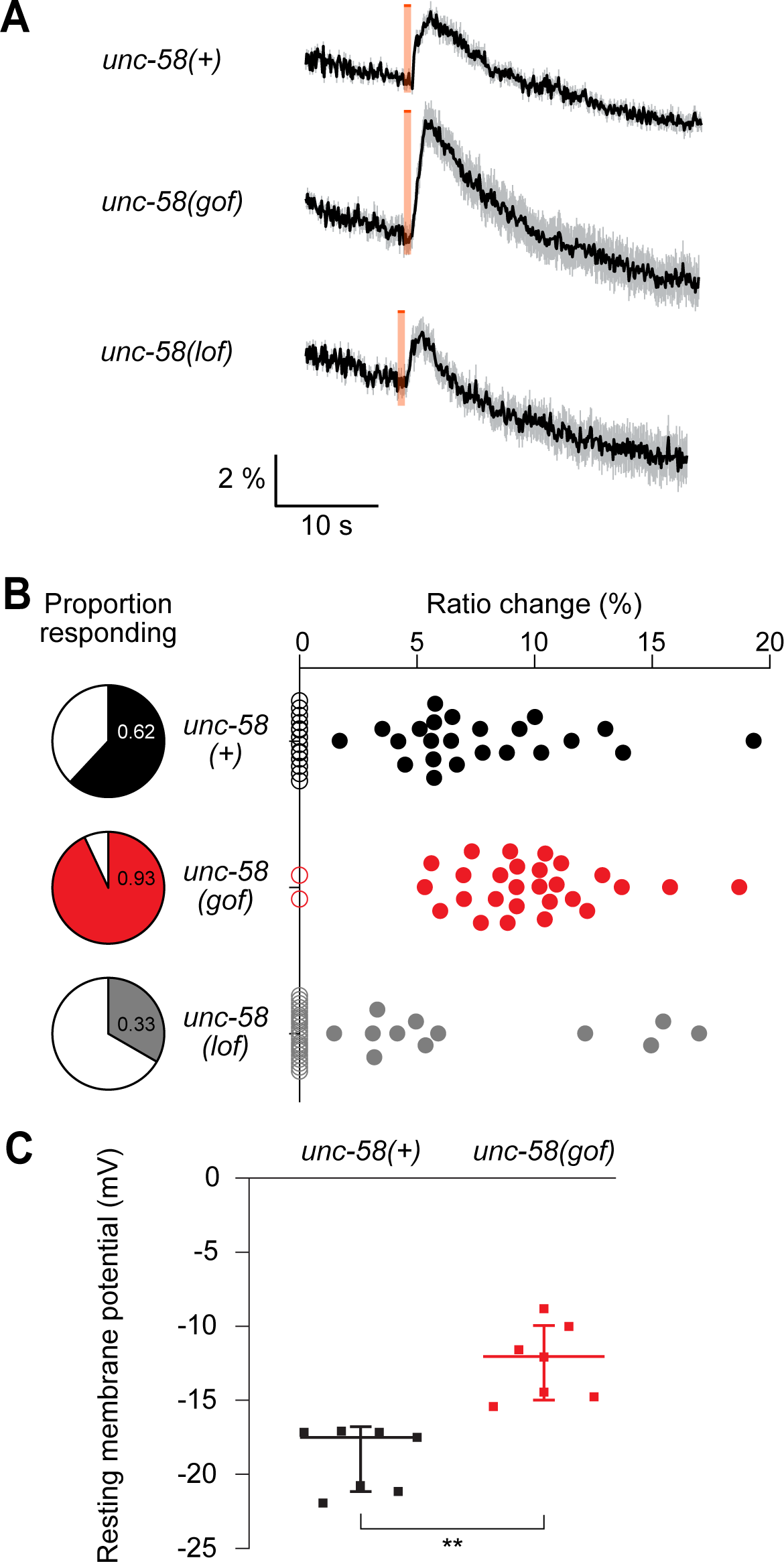
*unc-58* gain-of-function mutation increases neuronal and muscle excitability. **A** Average traces of Ca^2+^ responses of ALM neurons stimulated using a “short press” touch protocol in control (n=37), gain-of-function *unc-58(e665)* (n=29) and loss-of-function *unc-58(bln223)* (n=36) animals expressing the ratiometric calcium sensor YC3.60 in mechanosensory neurons. Orange bars indicate duration of physical stimulus. Black trace, mean CFP/YFP ratio change, ΔR (%). Grey trace, standard deviation. **B** Touch response of ALM mechanosensory neurons is modulated by mutation of *unc-58*. Left, proportion of ALM neurons responding to short press mechanical stimulus in wild type (n=37), gain-of-function mutant *unc-58(e665)* (n=29) and loss-of-function mutant *unc-58(bln223)* (n=36). P=0.0039 (wild type vs. *unc-58(e665))* and 0.0193 (wild type vs. *unc-58(bln223))*, Fisher’s exact test. Right, ratio change, ΔR (%) of calcium response. P<0.0001 and p=0.0084, respectively, Mann-Whitney test. Each data point represents the response of one ALM neuron. Non-responders, empty circles. **C** Resting membrane potential of *C. elegans* body wall muscles from wild type (n=7) and gain-of-function mutant *unc-58(bln205)* (n=7). Line, median; whiskers, standard deviation. Mann-Whitney test, ** p < 0.005.

We then tested the functional impact of *unc-58(gof)* in striated muscle cells. We recorded the resting membrane potential of body wall muscles using the whole-cell configuration of the patch-clamp technique. Instead of the hyperpolarization that would be expected for a hyperactive K^+^-selective K2P channel, we observed a significant depolarization of the muscle membrane in the *unc-58* gain-of-function mutant (Figure 2C).

### UNC-58 is a K2P channel permeable to sodium in physiological conditions

To directly investigate the ion selectivity of the UNC-58 channel, we used heterologous expression in *Xenopus laevis* oocytes. Expression of wild-type channels did not produce any detectable current when compared to non-injected oocytes (Figure 3A). However, when we recorded currents from oocytes expressing the UNC-58 L428F gain-of-function mutant, we observed a negative inward current whose intensity increased from −10 mV to −100 mV (Figure 3A). In addition, UNC-58 L428F expression resulted in a significant depolarization of the resting membrane potential (E_rev_= −15 ± 4.3 mV, median ± SD, n=6) compared to non-injected oocytes (E_rev_= −30 ± 7.3 mV, median ± SD, n=12, Kruskal-Wallis with Dunn’s post-hoc test, p<0.05), that is consistent with a depolarizing current (Supplementary Table ST2). As expected from the phenotypes of *unc-58(gof)* mutant animals, the current flowing through UNC-58 L428F is not consistent with UNC-58 being a K^+^ channel. Indeed, at this range of potentials one would expect a positive outward current above the K^+^ equilibrium potential (around −80 mV in physiological solutions) and a shift of the resting membrane potential towards the K^+^ equilibrium potential. We attempted to inhibit currents flowing through UNC-58 in order to isolate the fraction of the recorded current specific to UNC-58. We used the putative UNC-58 inhibitor endosulfan (Kasap and Dwyer, 2021), but did not observe any effect of this drug on the current (Figure S2).

**Figure 3.**
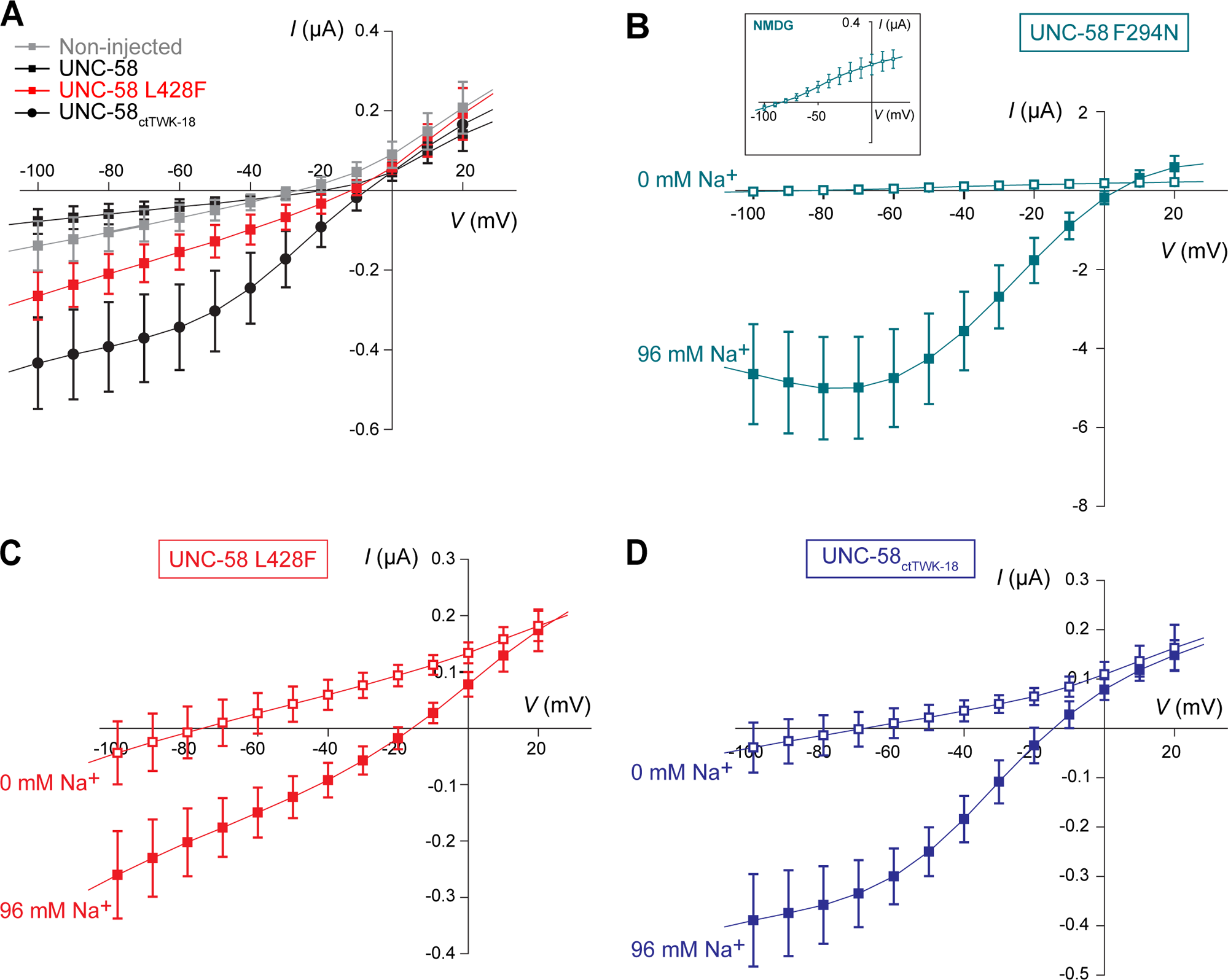
UNC-58 is an unconventional sodium-permeable K2P channel. **A** Current-voltage relationships obtained from *X. laevis* oocytes after injection of cRNA encoding wild-type (UNC-58, n=12), gain-of-function mutant (UNC-58 L428F, n=6), chimeric UNC-58 channels (UNC-58_ctTWK-18_, n=15), and non-injected oocytes (n=12). **B** Current-voltage relationships from *X. laevis* oocytes expressing UNC-58 F294N (n=5) in physiological extracellular solution (96 mM Na^+^, solid cyan square) or after ionic substitution of extracellular sodium by NMDG (0 mM Na^+^, open cyan square). Inset shows NMDG condition at a reduced scale. **C** Current-voltage relationships from *X. laevis* oocytes expressing UNC-58 L428F (n=7) in physiological solution (96 mM Na^+^, solid red square) or after ionic substitution of extracellular sodium by NMDG (0 mM Na^+^, open red square). **D** Current-voltage relationships from *X. laevis* oocytes expressing the UNC-58_ctTWK-18_ chimera (n=9) in physiological extracellular solution (96 mM Na^+^, solid blue square) or after ionic substitution of extracellular sodium by NMDG (0 mM Na^+^, open blue square). Each point represents the mean ± standard deviation. Curves were drawn for illustrative purposes only.

The unusual current recorded through UNC-58 L428F channels could be due to the effects of the gain-of-function mutation we introduced at position 428. We therefore attempted to obtain currents from UNC-58 channels harboring the wild-type leucine at this position. First, in an attempt to increase expression of the UNC-58 channel, we replaced the C-terminal cytoplasmic domain of the wild-type UNC-58 protein with the corresponding portion of TWK-18, a K^+^-selective *C. elegans* K2P channel that can be readily expressed in *Xenopus* oocytes (Kunkel et al., 2000). Expression of chimeric UNC-58_ctTWK-18_ channels elicited clear inward currents in the range of −100 to −10 mV and led to a significant depolarization of the resting membrane potential (E_rev_= −8 ± 4.0 mV median ± SD, n=15, Kruskal-Wallis with Dunn’s post-hoc test, p<0.0001, when compared with non-injected oocytes) (Figure 3A and Supplementary Table ST2). Next, we engineered a different gain-of-function mutation that targets a key residue in the second transmembrane domain, which we have shown to be a universal activating mutation for vertebrate and invertebrate K2P channels (Ben Soussia et al., 2019). We expressed this UNC-58 F294N mutant in *Xenopus* oocytes and again observed a rightward shift of the mean reversal potential and over 10-fold larger inward currents (Figure 3B).

To define the ionic nature of the inward current, we performed cation substitution experiments by replacing extracellular Na^+^ with the impermeable cation N-methyl-D-glucamine (NMDG). In the absence of extracellular Na^+^ (0 mM Na^+^ condition), we observed a dramatic reduction of the inward current accompanied by a marked leftward shift of the reversal potential towards hyperpolarized membrane potential values for all UNC-58 variants (Figure 3B, C, D, and Supplementary Table ST2). This clear shift of the reversal potential towards the equilibrium potential of potassium indicates that K^+^ ions likely flow through UNC-58 in these conditions.

Taken together these data provide compelling evidence that UNC-58 is a K2P channel that is constitutively permeable to Na^+^ in physiological conditions.

### UNC-58 contains an unusual cysteine in the SF1 selectivity filter

To determine the molecular basis of the unconventional selectivity of UNC-58, we aligned the two pore domains of UNC-58 with the sequences of several other K2P channels (Figure 4A and S3). We noticed the presence of a cysteine, C266, within the SF1 selectivity filter (T<u>C</u>GYG). Remarkably, among more than 60 K2P channel sequences from invertebrates and vertebrates, no other channel subunit harbors a cysteine residue at this position. Furthermore, we were unable to identify cysteine residues in the selectivity filter sequences of any other class of K^+^ channels, including voltage-gated and inwardly rectifying potassium channels.

**Figure 4.**
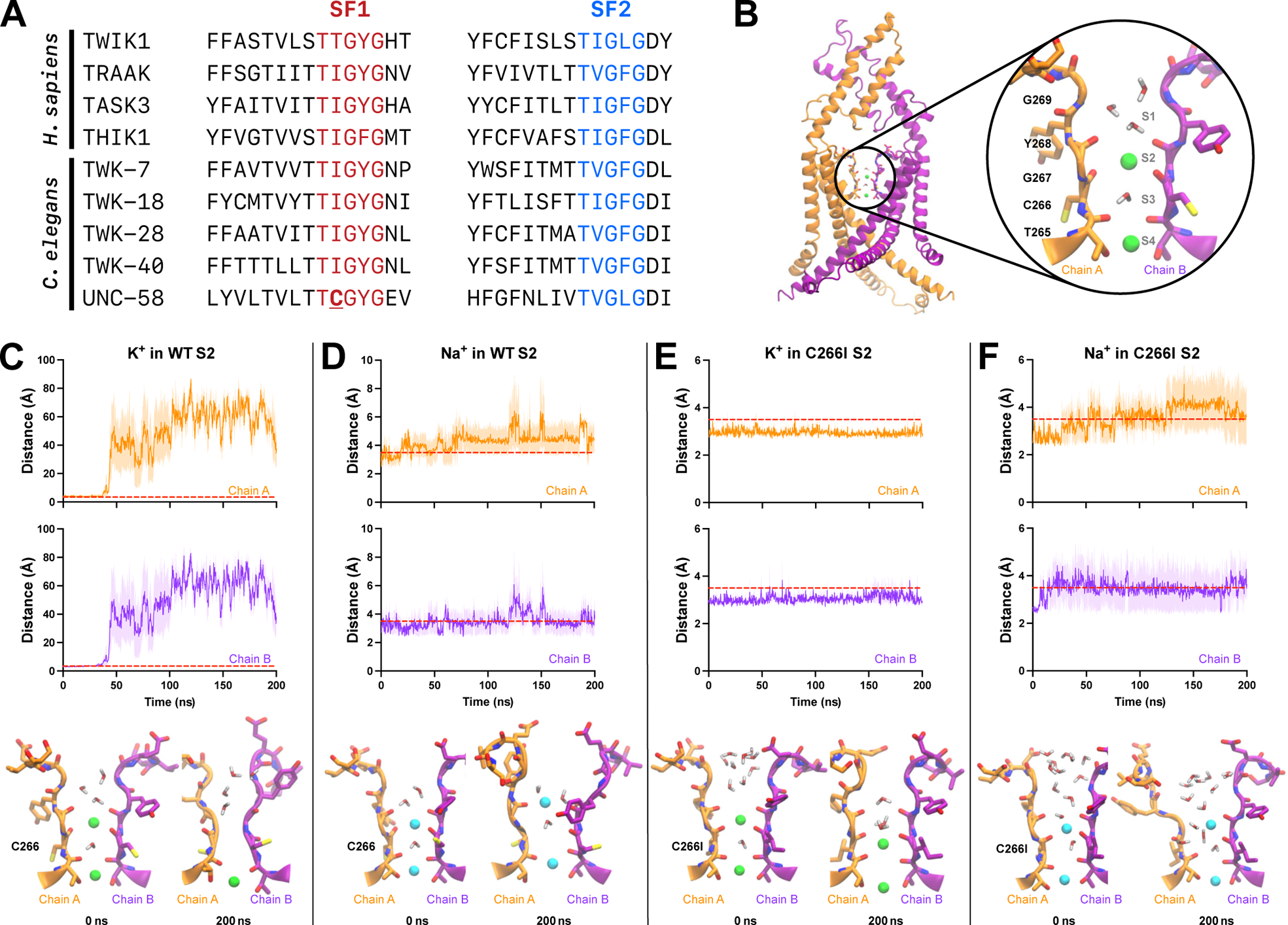
Ion coordination at the selectivity filter of UNC-58 channel. **A** Alignment of SF1 and SF2 selectivity filter sequences from selected human and *C. elegans* K2P channels. The SF1 and SF2 residues are labeled in red and blue, respectively. UNC-58 is the only channel containing a cysteine residue (in bold and underlined) at the second position of the TxGYG SF1 motif. **B** Structural model of the UNC-58 dimer, modeled with AlphaFold and used for molecular dynamics simulations. Individual subunits are labeled in orange and purple. Transmembrane helix M3 and SF2 have been omitted for clarity. The inset represents the SF1 region and shows opposing selectivity filter loops belonging two each subunit. K^+^ ions (green spheres) are coordinated at position S2 and S4 within the selectivity filter, while positions S1 and S3 are occupied by water molecules. **C-F** Upper panels indicate the distance between the S2 ion and the oxygen atom of the C266 carbonyl backbone during molecular dynamics (MD) simulations for wild type (WT) UNC-58 with K^+^ (C) or Na^+^ (D) and for the UNC-58 C266I mutant with K^+^ (E) or Na^+^ (F). Oxygen atoms within a confinement radius of 3.5 Å around the ion (red dotted lines) are counted as coordinating. Lower panels show representative licorice structures at the beginning (0 ns) and at the end (200 ns) of MD simulation. The residues of the SF1 selectivity filter of chain A and B are shown in orange and purple, respectively. K^+^ and Na^+^ ions are showed as green and cyan spheres, respectively.

To test whether this cysteine in SF1 could explain the unusual selectivity of UNC-58, we performed molecular dynamics (MD) simulations. In the absence of a crystal structure for UNC-58, we used the model deposited in the AlphaFold database as a basis for the 3D structure (Jumper et al., 2021). Based on this model, we observed that cysteine 266 could be involved in ion coordination at the S2 selectivity filter site (Figure 4B). We first ran three replicates of 200 ns-long MD simulations by placing either a K^+^ or a Na^+^ ion at the S2 position. To monitor ion coordination, we measured the distance of K^+^ or Na^+^ to the oxygen of the carbonyl backbone of C266 (O-C266) throughout the MD simulation. The carbonyl-K^+^ distance exhibited a more than tenfold increase, whereas the carbonyl-Na^+^ distance increased less than twice (Figure 4C, D) at the end of the simulations. This suggests that potassium was not efficiently coordinated in the UNC-58 selectivity filter.

We then calculated the occupancy of the ion placed initially at the S2 position, taking into account the time that the ion spends at a distance below the 3.5 Å threshold from O-C266 over the entire time of the MD simulations and expressed this occupancy in %. We found that O-C266 of both chains coordinated K^+^ with ∼13% of occupancy in UNC-58, and only at the beginning of the simulation (Figure 4C and Supplementary Table ST3). In contrast, Na^+^ ions were coordinated with ∼46% of occupancy (Figure 4D and Supplementary Table ST3).

Next, we measured the dihedral angle of C266 between the carbonyl oxygen and the nitrogen of the backbone as an additional indicator of ion coordination (Figure S4A, B). Compared to the beginning of the simulation, the dihedral angle changed by approximately 180° after 50 to 100 ns in the presence of K^+^ (Figure S4B). This change was observed in 2 out of 3 replicates for both subunits, and for one subunit in the last replicate. The consequence of this flipping motion was that O-C266 was no longer facing the pore, and was instead pointing to the opposite side (Figure S4A). Consistently, we observed that K^+^ was not retained in the selectivity filter (Figure 4C). In the presence of Na^+^, O-C266 flipped by 90° in only one subunit in each of the three replicates (Figure S4B). Na^+^ remained in the selectivity filter in all instances (Figure 4D).

These findings suggest that Na^+^ ions may be more effectively coordinated at the selectivity filter compared to K^+^ in the UNC-58 channel, consistent with the observed Na^+^ permeability in electrophysiological experiments. Additionally, these results provide evidence supporting the notion that C266 from the SF1 selectivity filter likely plays a central role in this alternative permeability.

### The cysteine residue in SF1 contributes to the sodium permeability of UNC-58

To assess the contribution of C266 to the unusual selectivity of UNC-58 channels, we substituted the C266 residue with a more conventional isoleucine in our model. We then conducted new molecular dynamics (MD) simulations on the resulting UNC-58 C266I channel.

Remarkably, the oxygen of the carbonyl backbone of C266I (O-C266I) coordinated K^+^ ion in the selectivity filter throughout the entire MD simulation with an occupancy of ∼94 % (Figure 4E and Supplementary Table ST3). Furthermore, the O-C266I/K^+^ distance remained below the 3.5 Å threshold in the three MD replicates, and the C266I dihedral angle remained constant at ∼155° throughout the simulation (Figure 4E and S4C, D). The mutant UNC-58 C266I also coordinated Na^+^ but in a more unstable manner, with an occupancy of ∼57 % (Figure 4F and Supplementary Table ST3). The O-C266I/Na^+^ distance increased during the simulation but remained close to that observed for the wild-type UNC-58 selectivity filter (Figure 4F). Finally, O-C266I flipped out in the presence of Na^+^ for one subunit, in two MD replicates, while it stayed stable in the remaining one (Figure S4D). In the three replicates, Na^+^ remained at the selectivity filter (Figure 4F). In conclusion, MD simulations predict that Na^+^ is more stable than potassium in the selectivity filter of wild-type UNC-58 channels, while mutation of the SF1 C266 to a more conventional isoleucine residue can dramatically increase potassium coordination.

To test the predictions of these molecular dynamics simulations, we engineered a C266I mutant in the context of the gain-of-function UNC-58 F294N and expressed this mutant channel in *Xenopus* oocytes. Functional characterization of the UNC-58 C266I F294N double mutant showed that the mean reversal potential of the current was significantly more negative for UNC-58 C266I F294N (E_rev_= −20 ± 8.7 mV, median ± SD, n=16) than for UNC-58 F294N (E_rev_= −2 ± 2.8 mV, median ± SD, n = 16) (Figure 5A, B). This result was consistent with a decrease in sodium selectivity of UNC-58 C266I F294N. We also noticed a drastic change in the shape of the current-voltage relationship compared the UNC-58 F294N single mutant, with the appearance of a strong inward rectification (Figure 5A). When extracellular Na^+^ was replaced by the impermeable cation choline, the current through UNC-58 C266I F294N was decreased (Figure 5C). For both UNC-58 F294N and UNC-58 C266I F294N, we observed a leftward shift of the reversal potential towards hyperpolarized membrane potentials, while Na^+^ substitution had no effect in water-injected oocytes (Figure S5). Notably, the shift was less pronounced in UNC-58 C266I F294N (−24 mV ± 8.5 mV) than in UNC-58 F294N (−61 mV ± 6.2 mV) (Figure 5D), suggesting that the C266I mutant is less permeable to Na^+^.

**Figure 5.**
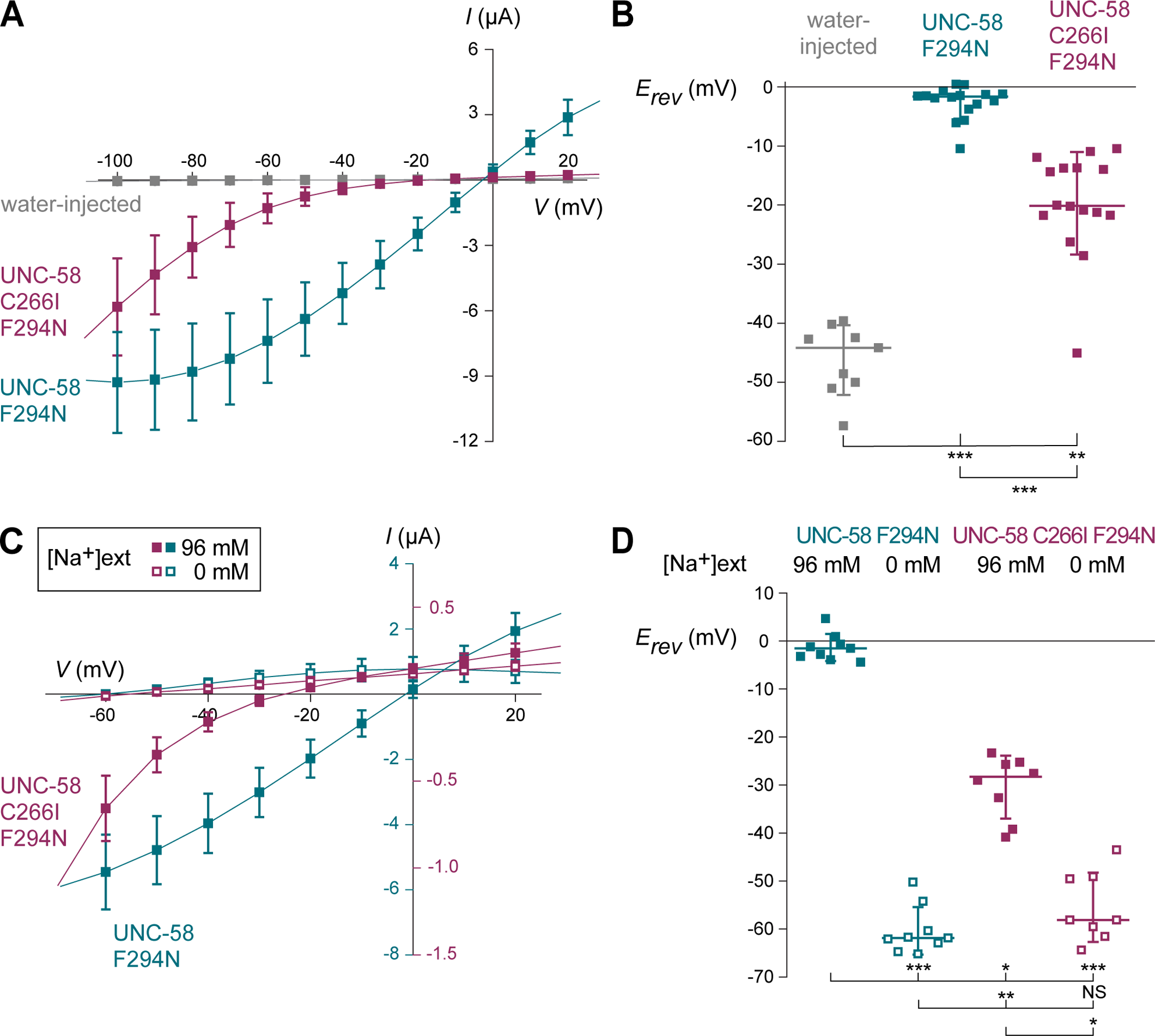
Cysteine 266 contributes to the sodium permeability of UNC-58. **A** Current-voltage relationship obtained from *X. laevis* oocytes injected with water (gray square, n = 9) or cRNA encoding UNC-58 F294N (cyan square, n = 16) or UNC-58 C266I F294N channels (burgundy square, n=16). Each point represents the mean ± standard deviation. Curves were drawn for illustrative purposes only. **B** Reversal potential of the current recorded from *X. laevis* oocytes expressing UNC-58 F294N and UNC-58 C266I F294N mutant channels. Current-voltage-relationships in A were fitted with a linear fit from −60 to −10 mV for oocytes injected by water (gray square), from −40 to +10 mV for oocytes expressing UNC-58 F294N (cyan square), and from −20 to +20 mV for oocytes expressing UNC-58 C266I F294N (burgundy square). Line, median; whiskers, standard deviation. Kruskal-Wallis test, p = 0.0031, followed by Dunn’s post-hoc test (** p < 0.005, *** p < 0.0005). **C** Current-voltage relationships obtained in *X. laevis* oocytes expressing UNC-58 F294N (cyan square, n = 16) or UNC-58 C266I F294N channels (burgundy square, n=16) in physiological extracellular solution (96 mM Na^+^, solid square) or after ionic substitution of extracellular sodium by choline (0 mM Na^+^, open square). Current values for oocytes expressing UNC-58 F294N should be read with the scale on the left of y-axis, current values for oocytes expressing UNC-58 C266I F294N should be read with the scale on the right of y-axis. Each point represents the mean ± standard deviation. Curves were drawn for illustrative purposes only. **D** Reversal potential of the current recorded from *X. laevis* oocytes expressing UNC-58 F294N and UNC-58 C266I F294N mutant channels. Current-voltage relationships in C were fitted with a linear fit from −40 to +10 mV for oocytes expressing UNC-58 F294N in physiological solution (solid cyan square), from −60 to −20 mV for oocytes expressing UNC-58 F294N in 0 mM Na^+^ solution (open cyan square), from −20 to 20 mV for oocytes expressing UNC-58 C266I F294N in physiological solution (solid burgundy square), and from −60 to −20 mV for oocytes expressing UNC-58 C266I F294N in 0 mM Na^+^ solution (open burgundy square). Line, median; whiskers, standard deviation. Kruskal-Wallis test, p<0.0001, followed by Dunn’s post-tests (NS: non-significant, * p<0.05, ** p<0.005, *** p<0.0005).

Altogether, these data provide compelling evidence that the highly unusual cysteine residue found in the SF1 selectivity filter contributes to the Na^+^ permeability of the UNC-58 channel.

## DISCUSSION

Here we report the first example of an evolutionary variation in the selectivity filter sequence of a two-pore domain (K2P) potassium channel that results in a dramatic difference in ion selectivity under physiological conditions. The *C. elegans* K2P channel UNC-58 has long been hypothesized to have an altered selectivity based on the hypercontracted phenotype of gain-of-function (*gof*) mutations (Salkoff, 2006). Indeed, when expressed in muscle and/or motor neurons, hyperactive K^+^ channels are expected to reduce membrane excitability and cause muscle relaxation. For example, flaccid paralysis is observed in *gof* mutants of the muscle TWK-18 (Kunkel et al., 2000) or the neuronal TWK-40 channel (Zhou et al., 2022). As expected, *gof* mutations in TWK-18 or TWK-40 give rise to an increased outward current and a decrease of the resting membrane potential (Ben Soussia et al., 2019; Yue et al., 2022). In contrast, we have shown here that UNC-58 carries an inward current and that hyperactivation of UNC-58 increases the resting membrane potential of muscle cells *in vivo*. This depolarization is reminiscent of the ones observed in *gof* mutants of the excitatory Ca^2+^ channel EGL-19 or the Na^+^ channel UNC-105 (Jospin et al., 2004, 2002). We cannot formally exclude the possibility that the muscle depolarization observed in *unc-58(gof)* mutants could be an indirect effect of the hyperactivation of motor neurons expressing UNC-58*(gof)* channels. However, degradation of hyperactive UNC-58 in muscle reduces the paralysis of *unc-58(gof)* animals, suggesting that UNC-58 does indeed play a role in muscle cells.

Although most of our experiments were performed on *unc-58(gof)* mutants, we also provide evidence that wild-type UNC-58 functions as a depolarizing channel under physiological conditions. Firstly, when we promoted the trafficking of UNC-58 in *Xenopus* oocytes by replacing its C-terminal region with that of the potassium-selective TWK-18 channel, we still observed an inward depolarizing current, similar to the one produced by UNC-58 *gof* channels. Secondly, the Ca^2+^ imaging experiments performed in the ALM mechanosensory neuron clearly demonstrate that neuronal excitability is decreased in the absence of UNC-58. These *in vivo* and *ex vivo* data indicate that the wild-type UNC-58 channel is indeed a depolarizing, Na^+^-conducting ion channel under physiological conditions. Most likely, gain-of-function mutations increase channel activity and therefore exacerbate this excitatory effect.

Our systematic analysis of K2P channel selectivity filter sequences revealed the presence of a highly unusual residue in the UNC-58 selectivity filter sequence (Figure S3). Indeed, the second position of the TxGYG motif harbors a cysteine, which is a unique feature in K2P channels since this position is usually occupied by an isoleucine, a valine or a threonine. Taken together, our results demonstrate that this residue contributes to the Na^+^ permeability of UNC-58. First, the reversal potential of the current carried by UNC-58 C266I mutant channels was shifted towards more negative potentials compared to that of wild-type UNC-58. Second, the substitution of Na^+^ by non-permeant ions induced a more pronounced shift of the reversal potential for wild-type UNC-58 compared to UNC-58 C266I. Finally, our MD simulations provided evidence for a higher permeability of K^+^ in the C266I channel than in wild-type UNC-58. However, it is difficult to conclude that UNC-58 C266I is a purely potassium-selective channel considering the reversal potential of the current carried by UNC-58 C266I. Being able to isolate UNC-58 currents from the endogenous currents would be necessary to further investigate the relative permeability to Na^+^ and K^+^, but we did not manage to selectively inhibit UNC-58 currents. Endosulfan has been proposed to block UNC-58 based on the improvement in locomotion observed in *unc-58(gof)* mutant animals after exposure to this compound (Kasap and Dwyer, 2021). However, even though we reproduced this effect on *unc-58(gof)* mutant animals (not shown), endosulfan did not alter UNC-58 currents recorded in *Xenopus laevis* oocytes. These results suggest that endosulfan does not act directly on UNC-58 but might modulate it by acting on a regulatory protein, or on other ion channels that may counteract the effect of increased UNC-58 activity.

While UNC-58 is the first K2P channel that has been shown to conduct Na^+^ in physiological conditions, some vertebrate K2P channels are known to lose K^+^ selectivity in particular conditions. TWIK1 selectivity relaxes at acidic pH or in low extracellular K^+^ concentration in *Xenopus* oocytes, cardiac muscle cell lines and human iPSC-derived cardiomyocytes (Chatelain et al., 2012; Chen et al., 2014; Liu et al., 2023; Ma et al., 2012, 2011). Interestingly, the SF1 sequence of TWIK1 harbors an unusual residue at the same position as the cysteine of UNC-58. In TWIK-1, the hydrophobic isoleucine found in most K2P SF1 selectivity filters (TIGYG) is replaced there by a threonine (Figure 4A and S3). Mutating this threonine residue to an isoleucine can restore TWIK1’s K^+^ selectivity at acidic pH or low extracellular K^+^ (Chatelain et al., 2012; Liu et al., 2023; Ma et al., 2011). Changes in K2P selectivity have also been reported in pathological conditions. A single point variant located at the equivalent position in the SF2 of TREK1 has been associated with ventricular tachycardia (Decher et al., 2017). In this pathological case, the SF2 sequence was mutated from TIGFG to T<u>T</u>GFG. This single point mutation drastically changes the channel’s selectivity to K^+^ ions, shifting the reversal potential by 40 mV towards more positive potentials. This behavior is consistent with the passage of Na^+^ through the modified selectivity filter, which was confirmed by molecular dynamics simulations (Decher et al., 2017). Yet, when equivalent mutations have been engineered in other potassium channels they did not alter channel selectivity. For instance, the voltage-dependent Shaker channel carrying a mutation equivalent to UNC-58 (T<u>C</u>GYG instead of T<u>V</u>GYG) did not conduct Na^+^ ions when expressed in *Xenopus* oocytes (Heginbotham et al., 1994).

Our MD simulations based on the AlphaFold model of UNC-58 offer insights into the possible underlying mechanisms that drive the non-selective behavior of this K2P channel. In the wild-type model, the carbonyl oxygens at position C266 (O-C266) of the two UNC-58 subunits rapidly flip out of the pore after the start of the simulation. This flipping leads to a loss of K^+^ coordination at the S2 site and the exit of the ion from the selectivity filter. Such a phenomenon has been observed during MD simulations for the TREK2 channel: flipping events of the selectivity filter carbonyl oxygens, especially at the S3 position, were observed more frequently in the non-conductive state, resulting in an intermittent closing of the selectivity filter, which prevented ions from passing through the filter (Brennecke and De Groot, 2018). Consistent with this observation, *in silico* free energy calculations in the KcsA channel suggest that the presence of a partially collapsed filter, even with just one flipped carbonyl group, is sufficient to disrupt potassium conduction and induce C-type inactivation (Bernèche and Roux, 2005). Interestingly, some K^+^ channels have been described to become more permeable to Na^+^ during C-type inactivation (Kiss et al., 1999; Starkus et al., 2000, 1997). When we ran the MD simulation in the presence of Na^+^ at the S2 site, we also observed that the carbonyl oxygen at position 266 flipped in WT UNC-58, as well as in the C266I model. It has been reported that the structural deformation in the selectivity filter caused by a carbonyl flipping significantly reduces the energy barriers for Na^+^ conduction in the prokaryotic non-selective NaK cation channels (Shi et al., 2018). Carbonyl flipping may therefore be important for Na^+^ flux through UNC-58: the unusual cysteine present in the UNC-58 selectivity filter, which drastically changes the lower coordination point of S2, could increase the energy barrier for K^+^ but decrease it for Na^+^.

The K2P UNC-58 thus behaves as a non-selective channel although its selectivity filter is close to that of other K^+^ channels. The non-selective hyperpolarization-activated cyclic nucleotide-gated (HCN) channel shares this feature: its selectivity filter (CIGYG) is close to that of K^+^ channels (TxGYG), yet HCN shows much lower selectivity for K^+^ than conventional potassium channels (see for review (Sartiani et al., 2017)). The crystal structure of HCN shows that the selectivity filter has a different shape compared to canonical K^+^ channels: the tyrosine side chain is rotated by 180 degrees compared to the tyrosine of the conventional K^+^ channel selectivity filter, and the carbonyl oxygen atoms from the tyrosine and the second glycine residues are not directed towards the ion pore and therefore cannot coordinate K^+^ at S1 and S2 (Lee and MacKinnon, 2017). The lack of coordination of K^+^ in the two outer binding sites of the selectivity filter has also been observed in another non-selective cation channel, the prokaryotic NaK channel (Gauss et al., 1998). It is thought that multi-occupancy of K^+^ ions in the selectivity filter is required for K^+^ permeation and that the absence of K^+^ coordination at S1 and S2 sites favors Na^+^ permeation (Bauer et al., 2022; Derebe et al., 2011; Lockless, 2015). Interestingly, the HCN channel has no ortholog in *C. elegans* (Hobert, 2013), whereas it is present in a wide range of vertebrate and invertebrate species (Sartiani et al., 2017). Given its excitatory activity, UNC-58 could partially fulfill the functions of an HCN channel, even though these channels carry currents with different biophysical properties, in particular with respect to membrane potential sensitivity. Indeed, wild-type UNC-58 currents do not exhibit the strong inward rectification characteristic of HCN channels. However, strikingly, the current-potential relationship observed for the UNC-58 C266I channel is very reminiscent of that recorded for HCN in invertebrates (Gauss et al., 1998).

The unusually large number of K2P channels encoded in the *C. elegans* genome could in principle provide the basis for a combinatorial code that would confer specific and diverse electrophysiological features to the limited set of *C. elegans* neurons. This hypothesis has been supported by systematic single-cell transcriptomic analyses as they reveal that distinct combinations of K2P channel genes are expressed in different cell types (Taylor et al., 2021). Remarkably, while some neuronal classes co-express over a dozen K2P channels, other neurons may express very few, and as little as a single K2P channel subunit. However, there is presently no direct way to predict the electrical behavior of a neuron from these gene expression profiles. In particular, since little is known about the ion selectivity or biophysical properties of most K2P channels in *C. elegans*, the contribution of each channel to neuronal excitability is unknown. Furthermore, K2P channels with divergent selectivity filter sequences may have unexpected functional properties, as we have demonstrated here for UNC-58. Thus, the potential for diversity in ion selectivity among *C. elegans* K2P channels may further enrich such a combinatorial ion channel code in a comparatively small and compact nervous system.

## ACKNOWLEDGEMENTS

We thank C. Frøkjær-Jensen, Wesley Hung, and Mei Zhen for plasmids, and Ithai Rabinowitch for constructing AQ1284. We thank Hannes Bülow, Bruno Allard and Jean-Louis Bessereau for comments and critical reading of the manuscript. Some strains were provided by the CGC, which is funded by NIH Office of Research Infrastructure Programs (P40 OD010440). I.B.S. was supported by AFM Téléthon (Alliance MyoNeurALP). This work was funded by the Agencia Nacional de Investigación y Desarrollo (ANID), grant Fondecyt Regular 1220656 (D.R.); by Fondecyt Regular 1230446 (W.G.); BQR, Bourse qualité recherche Université Lyon1 (O.A.) and a grant from the European Research Council (T.B., Ke*legans)* and Fondation Fyssen (T.B.); Medical Research Council MC-A023-5PB91 (W.R.S.); Wellcome Trust WT103784MA (W.R.S.) and Fonds voor Wetenschappelijk Onderzoek – Vlaanderen G079521N (W.R.S.).

## MATERIAL & METHODS

### *C. elegans* genetics and molecular biology

Except where stated, worms were raised at 20°C on nematode growth medium (NGM) and fed with *Escherichia coli* OP50 as previously described (Brenner, 1974). Strains and alleles used for this study are listed in Supplementary Table ST4 and ST5, respectively.

CRISPR/*Cas9* gene editing was performed using published protocols (Ben Soussia et al., 2019; Dokshin et al., 2018; El Mouridi et al., 2017). CRISPR/*Cas9-*induced molecular lesions were confirmed by Sanger sequencing for all alleles generated in this study. Germline transformation was performed by microinjection of plasmid DNA into the gonad of 1-day old *C. elegans* adult hermaphrodites.

Single-strand oligonucleotides, crRNA, and plasmids generated for this study are described in Supplementary Table ST5, ST6, and ST7, respectively.

### Identification of UNC-58 expression profile

Confocal imaging was performed using an inverted confocal microscope (Olympus IX83) equipped with a confocal scanner unit spinning-disk scan head (Yokogawa) and an EMCCD camera (iXon Ultra 888). Worms were imaged on 2% dry agar pads mounted in M9 solution containing 2% microbeads (Tebu-Bio, polystyrene polybeads 0.10 µm microsphere). Images in Figure S1 were acquired using a Zeiss confocal microscope (LSM880) with anesthetized worms (100 mM of sodium azide), mounted on 5% agarose pads on glass slides. Several z-stack images (each ∼0.45 mm thick) were acquired with the ZEN software. Representative images are shown following orthogonal projection of 2–10 z-planes. Image reconstruction was performed using ImageJ software.

The following neuronal markers were used to identify the neurons expressing *unc-58* by crossing the transcriptional reporter knock-in line *unc-58(bln259bln322*) with: *otIs388* (*eat-4^fosmid^::sl2::yfp::h2b*) (Serrano-Saiz et al., 2013), *otIs354* (*cho-1^fosmid^::sl2::yfp::h2b*) (Pereira et al., 2015), *otIs221* (*cat-1^prom^::gfp*) (Flames and Hobert, 2009), *nuIs1* (*glr-1^prom^::gfp*) (Hart et al., 1995), *evIs82b* (*unc-129^prom^::gfp*), *juIs76* (*unc-25^prom^::gfp*) (Huang et al., 2002), and *wdIs52* (*F49H12.4^prom^::gfp*).

### Locomotion test

Locomotion was assessed at 20°C on NGM plate seeded with a thin layer of OP50. Animal movements were recorded using an AZ100 Multizoom (Nikon) equipped with a Flash 4.0 CMOS camera (Hamamatsu Photonics) and analyzed using the *C. elegans* tracking software WormLab (MBF Bioscience). Measurements were performed for 50 s at 2 frames per second.

### *C. elegans* electrophysiology

Microdissection of *C. elegans* was performed as described previously (Lainé et al., 2011). Membrane potentials were recorded under current-clamp conditions in the whole-cell configuration using a MultiClamp 700B amplifier (Molecular Devices). Acquisition was done using the Clampex 10 software driving an Axon Digidata 1550 (Molecular Devices). Data were analyzed with Clampfit 10 (Molecular Devices) and graphed with Origin software (OriginLab). The bath solution contained (in mM) 150 NaCl, 5 KCl, 1 CaCl_2_, 1 MgCl_2_, 10 glucose, 15 HEPES, and sucrose to 330 mosm/L (pH 7.2), and the pipette solution (in mM) 120 KCl, 4 NaCl, 5 EGTA, 10 TES, 4 MgATP and sucrose to 320 mosm/L (pH 7.2). The resistance of recording pipettes ranged between 3 and 4 MΩ. All chemicals were obtained from Sigma-Aldrich. All experiments were performed at 20°C.

### Mechanosensory touch response assay

Worms were raised and assayed at 22 °C. Calcium imaging of anterior body touch stimulation of glued animals was essentially as described previously (Suzuki et al., 2003) except that a 1 second “short press” stimulus was used, in which a rounded capillary probe was displaced approximately 3 µm into the body. All stimulations were carried out level with the back of the terminal bulb of the pharynx, using day 1 adults. All animals expressed the cameleon YC3.60 under the control of *mec-4* promoter. Images were recorded at 10 Hz using an iXon EM camera (AndorTechnology), captured using IQ1.9 software (Andor Technology) and analyzed using a Matlab (MathWorks) program custom written by Ithai Rabinowitch (Rabinowitch et al., 2013). A rectangular region of interest (ROI) was drawn around the cell body and for each frame the ROI was moved according to the new position of the center of mass. Fluorescence intensity, F, was calculated as the difference between the sum of pixel intensities and the faintest 10% pixels (background) within the ROI. Fluorescence ratio R = F_yellow_/F_cyan_ (after correcting for bleed-through) was used for calculating the ratio change, expressed as a percentage of R_0_ (the average R within the first 3 seconds of recording).

### Two-electrode voltage-clamp electrophysiology in *Xenopus* oocytes

cDNA sequences were amplified using Phusion high-fidelity DNA polymerase (ThermoFisher Scientific) and assembled into the *Xenopus* oocyte expression vector pTB207 by 2- or 3-fragment Isothermal Ligation (Boulin et al., 2008). PCR-amplified sequences were validated by Sanger sequencing. The resulting vectors used in this study are listed in Supplementary Table ST7.

Capped mRNAs were synthesized *in vitro* from linearized expression vectors using the T7 mMessage mMachine kit (Ambion, Austin, TX, USA). Defolliculated *X. laevis* oocytes (Ecocyte Bioscience, Dortmund, Germany) were injected with 50 nL containing 30 ng of cRNA. Oocytes were kept at 18°C in ORII Calcium solution containing (in millimolar): 82.5 NaCl, 2 KCl, 1 MgCl_2_, 0.7 CaCl_2_, 5 HEPES, gentamicin (25 µg/mL), pH = 7.5 (with Trizma-Base).

Two-electrode voltage clamp (TEVC) experiments were performed 24 to 72 hours after microinjection. Oocytes were mounted in a small home-made recording chamber and continuously superfused with ND96 solution containing (in millimolar): 96 NaCl, 2 KCl, 1.8 CaCl_2_, 2 MgCl_2_, 5 HEPES. pH 7.4 was adjusted with Trizma base. For ion replacement experiments, Na^+^-free solution was prepared by replacing 96 mM NaCl by either 96 mM NMDG-Cl or 96 mM choline-Cl in the ND96 solution. Macroscopic currents were recorded using a Warner Instrument OC-725 amplifier, filtered at 10 kHz, digitized using a Digidata-1322 (Axon Instrument). For current visualization and stimulation protocol application, we used Axon pClamp 9 software (Molecular Devices, Sunnyvale, CA). Recording electrodes were pulled to 0.2-1.0 MΩ by using a horizontal puller (Sutter Instrument, Model P-97, USA) and filled with 3 M KCl. Currents were recorded in response to a voltage-step protocol consisting of a pre-pulse of −80 mV (80 ms duration) from a holding potential of −60 mV, followed by 10 mV steps (300 ms duration) from −150 mV to +40 mV, and return to a −60 mV holding potential. Current-voltage curves were obtained by plotting the steady-state currents at the end of each voltage step. Determination of reversal potentials was obtained by fitting the current trace with a linear equation in OriginPro.

### Molecular dynamics simulations (MDs)

To model K^+^/Na^+^ occupation in the UNC-58 selectivity filter we used the AlphaFold database model Q22271 (Varadi et al., 2022). The residues with pLDDT (per-residue confidence score) between 70 and 100 were kept (169 to 499). Then, the model was aligned with the TWIK1 crystal structure (PDB code: 3UKM) and both chains A and B ensembled. Four systems were built as follows: UNC-58 wild-type (WT) and K^+^, UNC-58 WT and Na^+^, UNC-58 C266I and K^+^, and UNC-58 C266I and Na^+^. K^+^ or Na^+^ were located at S2 and S4 positions and two water molecules were located at S1 and S3 (Egwolf and Roux, 2010). The models were prepared with the Protein Preparation Wizard software included in the Maestro Suite (Jacobson et al., 2002). Protonation states of amino acids were assigned at pH 7.0 with PROPKA (Olsson et al., 2011). The protein was embedded into a pre-equilibrated POPC bilayer and solvated using the SPC water model. The systems were neutralized with Cl^−^ or Na^+^/K^+^ ions and then, the ion concentration was set to 0.15 M KCl or NaCl. The Desmond membrane relaxation protocol (consisting of 6 stages) was used. After a proper system relaxation, two equilibrium MDs were performed in NPγT semi-isotropic ensemble. In the first one we applied a spring constant of 5 kcal×mol^−1^×Å^−2^ to the backbone atoms and 1 kcal×mol^−1^×Å^−2^ to the ions at selectivity filter for 10 ns. In the second one we applied a spring constant of 1 kcal×mol^−1^×Å^−2^ to the backbone atoms as well as the ions at selectivity filter for 10 ns. Both equilibrium MDs were performed with constant surface tension (0.0 bar×Å). Temperature and pressure were kept constant at 300 K and 1.01325 bar respectively by coupling to a Nose-Hoover Chain thermostat (Cheng and Merz, 1996) and Martyna-Tobias-Klein barostat (Martyna et al., 1994) with an integration time step of 2 fs. The simulations were performed with Desmond (Bowers et al., 2006) and the OPLS4 force field (Lu et al., 2021).

To run the production MDs, the last frame of the previous simulation was extracted and new MDs without restrains were run for 200 ns, the other parameters were conserved as previously described. Three replicas were performed *per* system, resulting in a total of 2400 ns MDs in this study. The MDs were analyzed with VMD (Humphrey et al., 1996) and Maestro tools.

