## Supplemental tables and figures for "Constitutive sodium permeability in a *C. elegans* two-pore domain potassium channel"

**Supplementary Figure S1. Expression pattern of *unc-58* in head and tail neurons.**

*unc-58* expression profile based on the transcriptional reporter *unc-58(bln259bln322 [unc-58::SL2::TagRFP-T])*. Cholinergic neurons are indicated in black, GABAergic neurons in blue, Glutamatergic neurons in yellow, Aminergic neurons in green and neurons with no identified neurotransmitter in red. GLR glial cells are indicated in pink. Each panel comes from a different confocal acquisition. Scale bars, 10 µm.

**Supplementary Movie SM1. Movement of *unc-58(e665)* gain-of-function mutants on solid media.**

Brightfield recording of mixed stage *unc-58(e665)* gain-of-function mutants on nematode growth media. 25 frames per second.

**Supplementary Movie SM2. Movement of *unc-58(bln205bln312)* gain-of-function mutants on solid media.**

Brightfield recording of young adult stage *unc-58(bln205bln312)* animals on nematode growth media. 2 frames per second.

**Supplementary Movie SM3. Movement of *unc-58(bln205bln312)* on solid media upon neuronal depletion of UNC-58.**

Brightfield recording of young adult stage *unc-58(bln205bln312); blnEx118* animals on nematode growth media. 2 frames per second.

**Supplementary Movie SM4. Movement of *unc-58(bln205bln312)* on solid media upon neuronal and muscle depletion of UNC-58.**

Brightfield recording of young adult stage *unc-58(bln205bln312); blnEx120* animals on nematode growth media. 2 frames per second.

**Supplementary Figure S2. Endosulfan has no direct effect on UNC-58 channels expressed in *Xenopus* oocytes.**

Current-voltage relationships obtained from *X. laevis* oocytes expressing UNC-58 L428F channels (n=6) in physiological extracellular solution (control, red square) or after 10 min incubation in 10 µM endosulfan (open gray circles).

Each point represents the mean ± standard deviation. Curves were drawn for illustrative purposes only.

**Supplementary Figure S3. Diversity of selectivity filter sequences in human, nematodes and flies.**

**A** Alignment of the SF1 and SF2 selectivity filter sequences of all human, *C. elegans*, and *Drosophila melanogaster* K2P channel subunits. The SF1 and SF2 residues are labeled in red and blue, respectively. UNC-58 is the only subunit containing a cysteine residue at the second position of the TxGYG selectivity filter motif.

**B** Sequence conservation along part of pore helix 1 and the SF1 selectivity filter computed from 66 vertebrate, insect, and nematode K2P channels, represented using WebLogo 3.

**C** Sequence conservation along part of pore helix 2 and the SF2 selectivity filter computed from 66 vertebrate, insect, and nematode K2P channels, represented using WebLogo 3.

**Supplementary Figure S4. Dihedral angles measured at residue 266 during molecular dynamics simulations.**

**A** Licorice representation of residues T265, C266, and G267 of wild type (WT) UNC-58 at the beginning (0 ns) and the end (200 ns) of MD simulation in the presence of $K^+$ at SF1 S2 position. The dihedral angle (encircled by a blue line) was measured between the oxygen and the nitrogen atoms of the C266 residue. The C266 carbonyl flipped during the simulation resulting in an increase of the dihedral angle of almost 180°.

**B** Evolution of dihedral angles for the C266 residue measured during the 200 ns MD simulations of WT UNC-58 subunit A (chain A, orange) and B (chain B, purple) in the presence of $K^+$ ($K^+_{S2}$ C266, upper panel) or $Na^+$ ($Na^+_{S2}$ C266, lower panel) at the S2 site. Three experimental replicates per system are displayed individually. Time is indicated as the radius of the circle.

**C** Licorice representation of residues T265, C266I, and G267 of the UNC-58 C266I mutant at the beginning (0 ns) and the end (200 ns) of MD simulation in the presence of $K^+$ at the S2 position. The dihedral angle (encircled by a blue line) was measured between the oxygen and the nitrogen atoms of the C266I residue. The C266I carbonyl oxygen did not flip during the simulation resulting in the maintenance of a constant dihedral angle.

**D** Evolution of dihedral angles for the C266I residue measured during the 200 ns MD simulations of UNC-58 C266I subunit A (chain A, orange) and B (chain B, purple) in the presence of $K^+$ ($K^+_{S2}$ C266I, upper panel) or $Na^+$ ($Na^+_{S2}$ C266I, lower panel) at the S2 site.

Three experimental replicates per system are displayed individually. Time is indicated as the radius of the circle.

**Supplementary Figure S5. Extracellular sodium substitution has no effect on the reversal potential of water-injected oocytes.**

**A** Current-voltage relationships obtained in *X. laevis* oocytes injected with water (n=10) in physiological solution (96 mM $Na^+$, solid gray square) or after ionic substitution of extracellular $Na^+$ by NMDG (0 mM $Na^+$, open gray square). Each point represents the mean ± standard deviation. Curves were drawn for illustrative purposes only.

**B** Mean reversal potential of *X. laevis* oocytes injected with water. Current-voltage-relationships were fitted with a linear fit from -60 to -10 mV in physiological solution (96 mM $Na^+$, solid square) or in 0 mM $Na^+$ solution (gray square). Line, median; whiskers, standard deviation. Wilcoxon test.

#### Supplementary materials

**Supplementary Table ST1.** Expression profile of *unc-58* in the nervous system

| Name | Neurotransmitter identity | Cell type | Transgene used for determination |
|---|---|---|---|
| AIN | ACh | Interneuron | *otIs354* |
| ALM | Glu | Sensory neuron | *otIs388* |
| ALN | ACh | Sensory neuron | *otIs354* |
| AS | ACh | Motor neuron | at the limit of detection with *otIs354* |
| ASK | Glu | Sensory neuron | *otIs388* |
| AUA | Glu | Interneuron | *otIs388* |
| AVH | Unknown (orphan) | Interneuron | by position, a pair next to RIV |
| AVK | Unknown (orphan) | Interneuron | by position, a pair next to AIY |
| DA | ACh | Motor neuron | *evIs82b / otIs354* |
| DB02-07 | ACh | Motor neuron | *evIs82b / otIs354* |
| FLP | Glu | Sensory neuron | *otIs388* |
| GLR | | Glial cells | by position |
| I4 | Unknown (orphan) | Polymodal neuron | by position next to AIN on left side |
| IL1 | Glu | Polymodal neuron | *otIs388* |
| M1 | ACh | Polymodal neuron | by position next to AIN on right side |
| M2 | ACh | Polymodal neuron | by position |
| M3 | Glu | Polymodal neuron | *otIs388* |
| OLL | Glu | Sensory neuron | *otIs388* |
| PDA | ACh | Motor neuron | *otIs354* |
| PDE | DA | Sensory neuron | by position a pair next to PVD |
| PHC | Glu | Sensory neuron | *otIs388* |
| PLM | Glu | Sensory neuron | *otIs388* |
| PLN | ACh | Sensory neuron | *otIs354* |
| PVD | Glu | Sensory neuron | *wdIs52 and otIs388* |
| PVR | Glu | Interneuron | *otIs388* |
| RIG | Glu | Interneuron | *otIs388* |
| RIH | ACh | Interneuron | *otIs354* and *otIs221* |
| RIM | Glu & Tyramine | Motor neuron | *otIs388 and nuIs1* |
| RIR | ACh | Interneuron | *otIs354* |
| RIV | ACh | Interneuron | *otIs354* |
| RME_DV | GABA | Motor neuron | *juIs76* |
| SAA | ACh | Interneuron | *otIs354* |
| SAB | ACh | Motor neuron | *otIs354* |
| SDQ | ACh | Interneuron | *otIs354* |
| SIB | ACh | Motor neuron | *otIs354* |
| SMB | ACh | Motor neuron | *otIs354* |
| URA | ACh | Sensory neuron | *otIs354* |
| URY | Glu | Sensory neuron | *otIs388* |
| VA2-12 | ACh | Motor neuron | *otIs354* |
| VB | ACh | Motor neuron | *otIs354* |
| VC4 | ACh | Motor neuron | *otIs354* |
| VD_DD | GABA | Motor neuron | *juIs76* |

**Supplementary Table ST2.** Reversal potentials ($E_{rev}$).

| | | $E_{rev}$ (mV) median ± SD | | Statistical significance (n) |
|---|---|---|---|---|
| Figure 3A | Non-injected | -30 ± 7.3 | | (12) |
| | UNC-58 | -23 ± 4.0 | | p = 0.59 (12) |
| | UNC-58 L428F | -15 ± 4.3 | | p = 0.03 (6) |
| | UNC-58$_{ctTWK-18}$ | -8 ± 3.9 | | p < 0.0001 (15) |
| | | with $Na^+$ | without $Na^+$ | |
| Figure 3B | UNC-58 F294N | 4 ± 2.1 | -82 ± 8.5 | p = 0.0625 (5) |
| Figure 3C | UNC-58 L428F | -18 ± 4.4 | -83 ± 24.3 | p = 0.015 (7) |
| Figure 3D | UNC-58$_{ctTWK-18}$ | -12 ± 5.0 | -82 ± 28.4 | p = 0.0039 (9) |
| Figure S5B | Non-Injected | -31 ± 6.2 | -34 ± 6.5 | p = 0.1309 (10) |

Wilcoxon matched-pairs signed rank test, except Kruskal-Wallis with Dunn's post-hoc test for data of Figure 3A.

**Supplementary Table ST3.** Ion occupancy.

| System | % of ion coordination | | |
|---|---|---|---|
| | Chain A | Chain B | Average |
| UNC-58 WT + $K^+$ | 9 ± 0.5 | 16 ± 0.7 | 13 ± 0.4 |
| UNC-58 WT + $Na^+$ | 37 ± 0.9 | 55 ± 0.9 | 46 ± 0.6 |
| UNC-58 C266I + $K^+$ | 96 ± 0.3 | 91 ± 0.5 | 94 ± 0.4 |
| UNC-58 C266I + $Na^+$ | 56 ± 0.9 | 57 ± 0.9 | 56 ± 0.5 |

Values are Average ± SEM of three replicas (200 ns each) per system.

**Supplementary Table ST4.** Strains used for this study.

| **Figure** | **Strain** | **Genotype** |
|---|---|---|
| 1D, S1 | JIP1444 | *unc-58(bln259bln322) X* |
| 1E | JIP1473 | *unc-58(bln323) X* |
| | JIP1474 | *unc-58(bln324) X* |
| SM1 | CB665 | *unc-58(e665) X* |
| 1F, 1G, SM2 | JIP1451 | *unc-58(bln205bln312) X* |
| 1F, 1G, SM3 | JIP1490 | *unc-58(bln205bln312) X; blnEx118* |
| 1F, 1G, SM4 | JIP1492 | *unc-58(bln205bln312) X; blnEx120* |
| 2A, B | AQ1284 | *ijIs130* |

| | | |
|---|---|---|
| | JIP1723 | *unc-58(e665) X; ijIs130* |
| | JIP1738 | *unc-58(bln223) X; ijIs130* |
| 2C | JIP1135 | *unc-58(bln205) X* |

**Supplementary Table ST5.** Alleles used for this study.

| **Allele** | **Description** | **Source** |
|---|---|---|
| *bln205* | UNC-58 L428F ; GTttcTCAGTAGTGACCATG ; using sgRNA pPT39 and repair template oPT73. | this study |
| *bln223* | *unc-58* loss of function allele; deletion from exon 1a to 10; Breakpoints AGGCCCACTATTTCAATCCAAGAAG//ATTCACTACTTTGGTCGAGCAAAAC; using sgRNAs pPT35, pPT44, pPT51, pPT53. | (Rawsthorne-Manning et al., 2022) |
| *bln259* | *unc-58::SL2::TagRFP-T* transcriptional reporter; using sgRNA pMD8 and repair template pPT89 in JIP1232. | this study |
| *bln312* | *unc-58::TagRFP-T::ZF1* ; using sgRNA pMD8 and repair template pPT81 in JIP1232. | this study |
| *bln322* | Reversion of *bln205* [L428F] to wild type [L428]; GTCTTTCCGTAGTGACtATGTGC ; using with sgRNA pPT35 and repair template pPT67 in JIP1355. | this study |
| *bln323* | *mNeonGreen* translational fusion to isoform b. | this study |
| *bln324* | *mNeonGreen* translational fusion to isoform a. | this study |
| *blnEx118* | pPT85 [*acr-2$^{prom}$::zif-1::SL2::ebfp*] 50 ng/µL; pCFJ421 [*myo-2$^{prom}$::gfp::h2b*] 10 ng/µL | this study |
| *blnEx120* | pPT83 [*myo-3$^{prom}$::zif-1::SL2::ebfp*] 20 ng/µL; pT85 [*acr-2$^{prom}$::zif-1::SL2::ebfp*] 20 ng/µL; pCFJ421 [*myo-2$^{prom}$::gfp::h2b*] 10 ng/µLpPT83 20 ng/µL; pT85 20 ng/µL; pCFJ421 10 ng/µL | this study |
| *e665* | UNC-58 L428F, ChrX:10105660 C/T | CGC |
| *evIs82b* | *unc-129$^{prom}$::gfp* | CGC |
| *ijIs130* | *mec-4$^{prom}$::yc3.60* | Ithai Rabinowitch, unpublished |
| *juIs76* | *unc-25$^{prom}$::gfp* | (Huang et al., 2002) |
| *nuIs1* | *glr-1$^{prom}$::gfp* | (Hart et al., 1995) |
| *otIs354* | *cho-1$^{fosmid}$::SL2::yfp::h2b* | (Pereira et al., 2015) |
| *otIs388* | *eat-4$^{fosmid}$::SL2::yfp::h2b* | (Serrano-Saiz et al., 2013) |

| | | |
|---|---|---|
| *otIs221* | *cat-1*$^{prom}$*::gfp* | (Flames and Hobert, 2009) |
| *wdIs52* | *F49H12.4*$^{prom}$*::gfp* | CGC |

**Supplementary Table S6.** Sequences of single-strand oligonucleotide repair templates, synthetic crRNA, and crRNA from pPT2-based single guide RNA expression vectors.

| **Name** | **Sequence (5' to 3')** | **Corresponding allele** |
|---|---|---|
| | *Single-strand oligonucleotide repair template (ssODN)* | |
| oPT73 | GGTACCTAATACGACTCACTATAGGGGTCGAAGTCTGG TGCATGATCC | *bln205* |
| | *sgRNA expression vectors* | |
| pMD8 | GCUACCAUAGGCACCACGAG | (D'Alessandro et al., 2018) |
| pPT35 | GGACGCAAGAUCCACGCACA | *bln223, bln322* |
| pPT39 | UCCACGCACAUGGUCACUA | *bln205* |
| pPT44 | UUCGUUCUCAAAAUUGUUUG | *bln223* |
| pPT51 | UGUCAGGUAAGAAGAACAAU | *bln223* |
| pPT53 | UCCACGCACAUGGUCACUA | *bln223* |

**Supplementary Table ST7.** *Xenopus* oocyte expression plasmids.

| **Figure** | **Name** | **Construct** | **Description** |
|---|---|---|---|
| 3A | pMM8 | UNC-58 | Wild-type cDNA. |
| 3A, C<br>S2 | pMM10 | UNC-58 L428F | Gain-of-function mutant corresponding to *e665.* |
| 3A, D | pIBS11 | UNC-58$_{ctTWK-18}$ | Replacement of UNC-58 C-terminus after A438 by TWK-18 C-terminus starting at Q302. |
| 3B<br>5A-D | pOA18 | UNC-58 F294N | TM2.6 gain-of-function mutant (Ben Soussia et al., 2019). |
| 5A-D | pOA20 | UNC-58 C266I F294N | TM2.6 gain-of-function and C266I mutant. |

### Figure S1

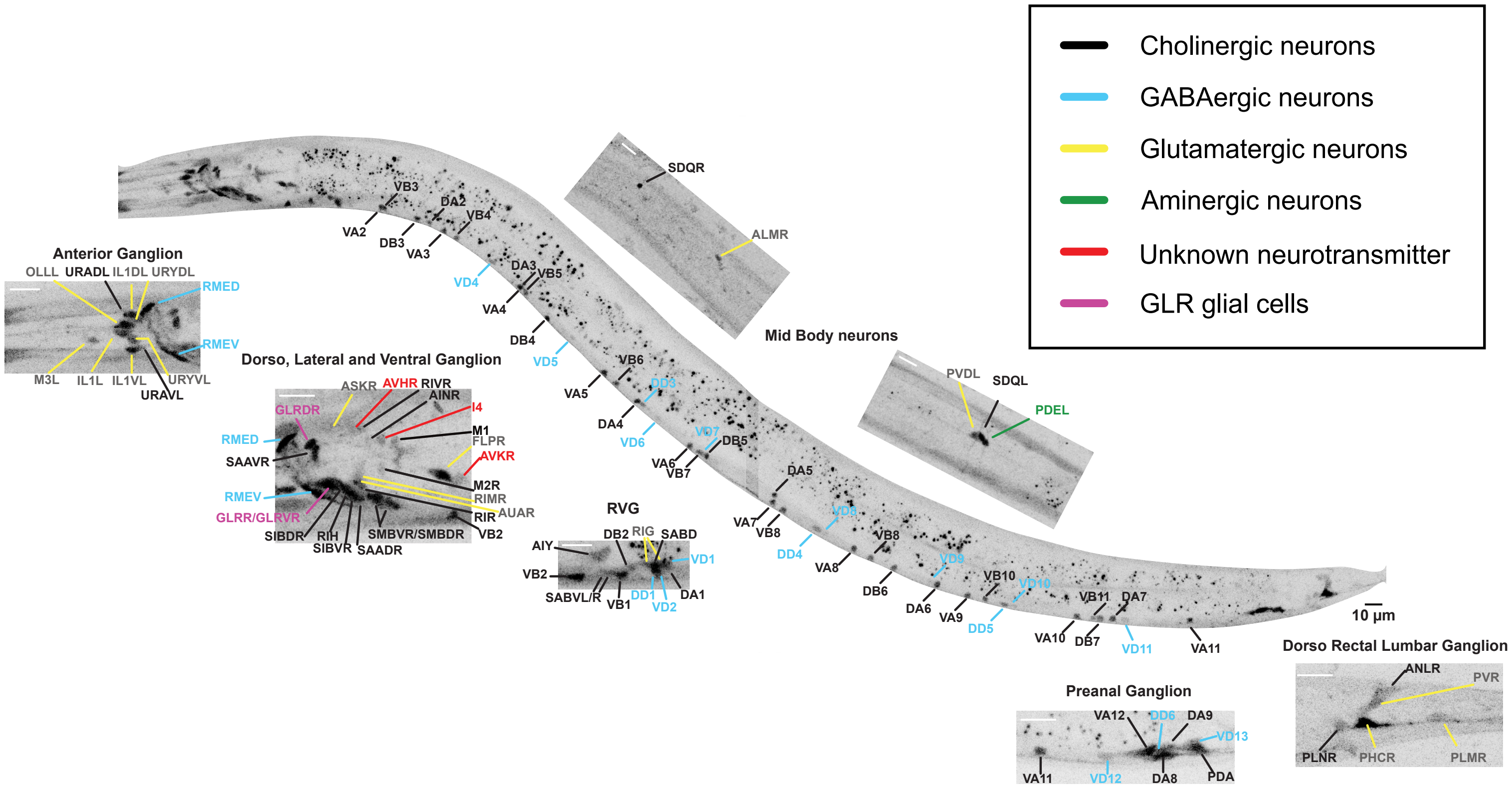

Cholinergic neurons
GABAergic neurons
Glutamatergic neurons
Aminergic neurons
Unknown neurotransmitter
GLR glial cells
Anterior Ganglion
OLLL
URADL
IL1DL
URYDL
RMED
RMEV
M3L
IL1L
IL1VL
URYVL
URAVL
Dorso, Lateral and Ventral Ganglion
ASKR
AVHR
RIVR
AINR
GLRDR
I4
M1
FLPR
AVKR
RMED
SAAVR
M2R
RIMR
RMEV
AUAR
GLRR/GLRVR
RIR
VB2
SIBDR
RIH
SIBVR
SMBVR/SMBDR
SAADR
RVG
AIY
DB2
RIG
SABD
VD1
VB2
SABVL/R
VB1
DD1
VD2
DA1
VB3
DA2
VB4
VA2
DB3
VA3
DA3
VB5
VD4
VA4
DB4
VD5
VB6
DD3
VA5
DA4
VD6
VD7
DB5
VA6
VB7
DA5
VA7
VB8
VD8
DD4
VB8
VA8
DB6
VD9
VB10
VD10
DA6
VA9
DD5
VB11
DA7
VA10
DB7
VD11
VA11
10 µm
SDQR
ALMR
Mid Body neurons
PVDL
SDQL
PDEL
Preanal Ganglion
VA12
DD6
DA9
VD13
VA11
VD12
DA8
PDA
Dorso Rectal Lumbar Ganglion
ANLR
PVR
PLNR
PHCR
PLMR

### Figure S2

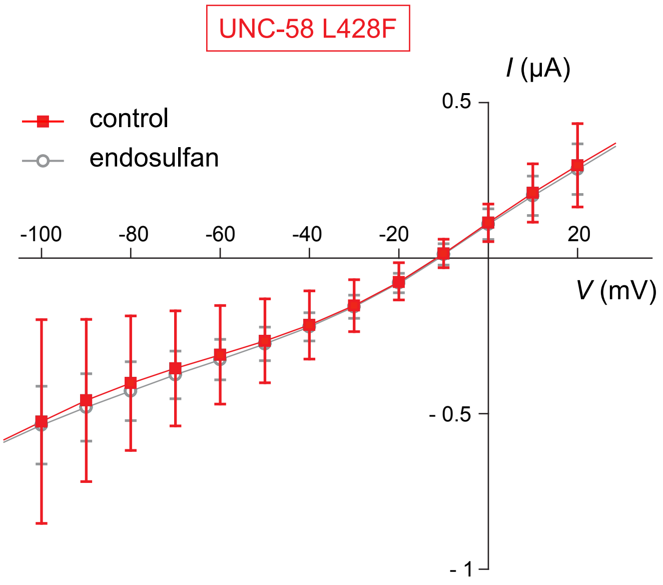
UNC-58 L428F
I (μA)
0.5
control
endosulfan
-100
-80
-60
-40
-20
20
V (mV)
- 0.5
- 1

### Figure S3

**A**

| | | SF1 | SF2 |
|---|---|---|---|
| *H. sapiens* | TWIK1 | FFASTVLSTTGYGHT | YFCFISLSTIGLGDY |
| | TWIK2 | FFASTLITTVGYGYT | YFCFISLSTIGLGDY |
| | TRAAK | FFSGTIITTIGYGNV | YFVIVTLTTVGFGDY |
| | TREK1 | FFAGTVITTIGFGNI | YFVVITLTTIGFGDY |
| | TREK2 | FFAGTVITTIGYGNI | YFVVVTLTTVGFGDF |
| | TASK1 | YFAITVITTIGYGHA | YYCFITLTTIGFGDY |
| | TASK3 | YFAITVITTIGYGHA | YYCFITLTTIGFGDY |
| | TASK5 | YFAITVITTIEYGHA | YYCFITLTTIGFGDF |
| | TASK2 | IFAATVITTIGYGNV | YYSFITISTIGFGDF |
| | TALK1 | FFAGTVVTTIGYGNL | YFAFITLSTIGFGDY |
| | TALK2 | FFSVSTITTIGYGNL | YFAFITLSTVGFGDY |
| | TRESK | FFCCTVFSTVGYGYI | YFCFVTLTTIGFGDT |
| | THIK1 | YFVGTVVSTIGFGMT | YFCFVAFSTIGFGDL |
| | THIK2 | YFVGTVVSTIGFGMT | YFCFVTFSTIGFGDL |
| | KCNK7 | LFAASILTTTGYGHM | YFCFSSLSTIGLEDL |
| *C. elegans* | EGL-23 | LFAFTVITTIGYGNV | YFNFVTLTSIGLGDI |
| | UNC-58 | LYVLTVLTTCGYGEV | HFGFNLIVTVGLGDI |
| | SUP-9 | YFATTVITTIGYGHS | YYCFVTLTTIGFGDY |
| | TWK-2 | FFAATTMATIGYGNI | YYSFISLTTIGFGDI |
| | TWK-3 | VFTTTTVIPVGYGYI | YFSLTSIFTIGFGDL |
| | TWK-4 | FFTTTVLTSIGYGNL | YFCLVSLLTVGFGDL |
| | TWK-6 | FFSWTLYSTVGYGSL | YFGIITMFLIGIGDI |
| | TWK-7 | FFAVTVVTTIGYGNP | YWSFITMTTVGFGDL |
| | TWK-8 | YFAGTLFTTIGYGDI | YFMFISLSTIGLGDV |
| | TWK-9 | IFAFTVITTIGYGHV | YFNFVTLTTIGLGDF |
| | TWK-10 | LFTATTIIPVGYGYI | YFSTITIFTIGFGDI |
| | TWK-11 | LYAGTIFTTIGYGNI | YFCFISLTTIGLGDV |
| | TWK-12 | FYAGQLYTTIGYGYP | YFFIVSISTVGLGDM |
| | TWK-13 | FFSWTAITTIGYGHI | YFCFVTVTTIGFGDI |
| | TWK-14 | FFSATVISTIGFGTS | YFCFISFATIGFGDY |
| | TWK-16 | LYGFGIVTTLGYNRI | YFNFLCLTAIDFGQL |
| | TWK-17 | LYALTVITSTGYDHV | FFSFNTITTIGLGNI |
| | TWK-18 | FYCMTVYTTIGYGNI | YFTLISFTTIGFGDI |
| | TWK-20 | YFATVVITTVGYGHS | YFCMITFSTIGFGDL |
| | TWK-21 | FFSATIFTTIGYGNL | YFFFVSLSTIGFGDI |
| | TWK-22 | FFSATIFTTIGYGNL | YFFFVSLSTIGFGDV |
| | TWK-23 | LFSFTILTTIGYGNV | YFSYMSLTTIGLGDI |
| | TWK-24 | YYSASLYTTIGYGNF | YFFFTSLTTIGLGDV |
| | TWK-25 | FFSMNVYTTTGYGSI | YFSFISMSTIGLGDV |
| | TWK-26 | FYSMTLFSTIGYGTI | YFTFISLTTIGLGDV |
| | TWK-28 | FFAATVITTIGYGNL | YFCFITMATVGFGDI |
| | TWK-29 | LYSLGILTTLGYGKI | YYAFICLTAIEYGDI |
| | TWK-30 | FFTTTMLTSIGYGYV | YFIFISILTVGFGDI |
| | TWK-31 | FYVGTLFTTIGYGNI | YFFCISLLTIGLGDI |
| | TWK-32 | FFAATTLTSIGYGID | YFVMMSVLTIGFGDL |
| | TWK-33 | FFSMNVYTTTGYGSI | YFSFISLSTIGLGDI |
| | TWK-34 | FFSMNVYTTTGYGSI | YFSFISLSTIGLGDI |
| | TWK-35 | LFAGTICTTIGYGHI | YFFFTSLSTVGLGDL |
| | TWK-36 | FFSMNVYTTTGYGSI | YFSFISLTTIGLGDV |
| | TWK-37 | AYVITCITTIGYGEL | YFSFSTFSTIGFGDF |
| | TWK-39 | LYSITVFTTIGYGHI | YYCFITLSTIGFGDI |
| | TWK-40 | FFTTTLLTTIGYGNL | YFSFITMTTVGFGDI |
| | TWK-42 | YYAGTIYTTIGYGDL | YFFFISMTTIGFGDF |
| | TWK-43 | LFCFTVITTIGYGNV | YFNFVTLTTIGLGDL |
| | TWK-44 | FLAVTTYTTIGYGNI | YFFFCSLTTIGYGDV |
| | TWK-45 | FYSMTLFSTIGYGTI | YFTFISLTTIGLGDV |
| | TWK-46 | FFAGTLISTVGYGRV | YYCFVSLTTIGLGDF |
| | TWK-48 | LYSLTVITTIGYGNT | YFCFISLTTIGFGDK |
| *D. melanogaster* | ORK1/KCNK0 | FFAFTVCSTVGYGNI | YYSYVTTTTIGFGDY |
| | dTask-6 | YYATTVLTTIGYGHS | YYCFITLTTIGFGDM |
| | dTask-7 | YFSTVVLAMIGYGHS | YYCFVTLTTIGFGDY |
| | sandman | LYSLTVITTIGYGNI | YFCLISLSSIGFGDL |
| | CG10864 | MFCLSVITMIGYGNM | YFCMTSLCKIGFGDF |
| | CG34396 | VYSATIYTTIGYGHI | YYVFISMSTIGFGDL |
| | CG43155 | FFSSTVLTTIGYGNI | YFCFITMTTIGFGDL |
| | Galene | FYSIIVITTIGYGHI | YFCFITLTTIGFGDF |

**B**

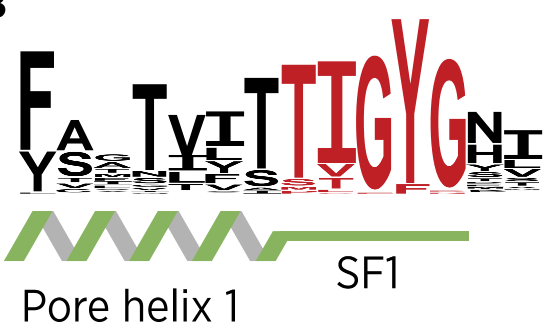


**C**

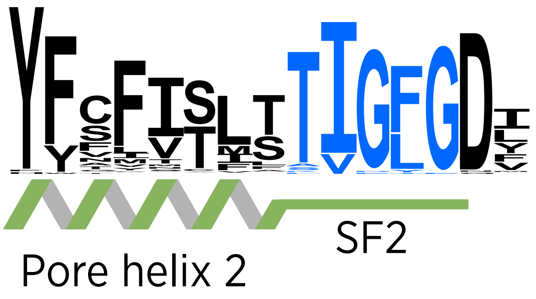

### Figure S4

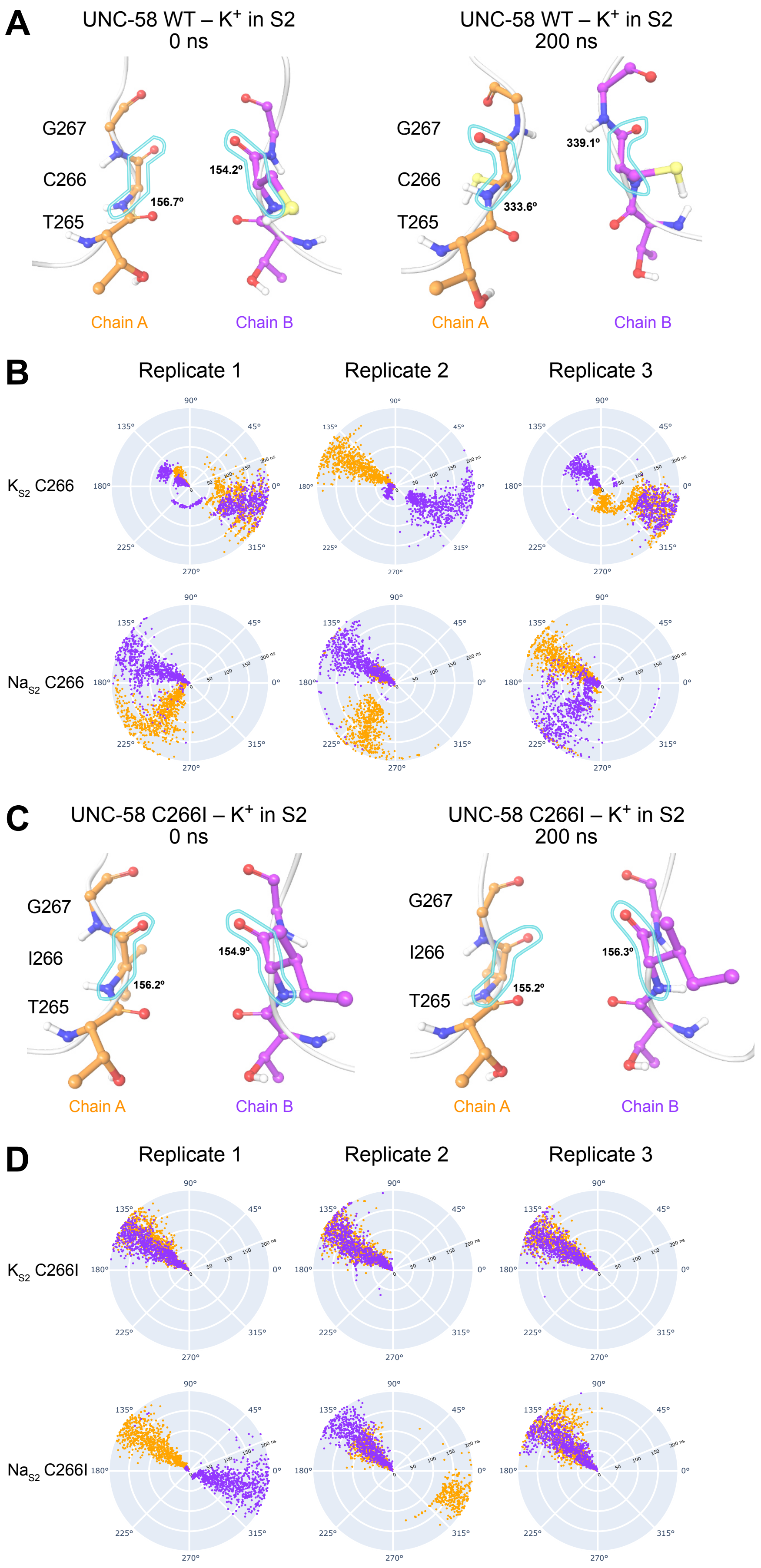
A
UNC-58 WT – K+ in S2
0 ns
UNC-58 WT – K+ in S2
200 ns
G267
C266
T265
156.7º
154.2º
333.6º
339.1º
Chain A
Chain B
Chain A
Chain B
B
Replicate 1
Replicate 2
Replicate 3
KS2 C266
NaS2 C266
C
UNC-58 C266I – K+ in S2
0 ns
UNC-58 C266I – K+ in S2
200 ns
G267
I266
T265
156.2º
154.9º
155.2º
156.3º
Chain A
Chain B
Chain A
Chain B
D
Replicate 1
Replicate 2
Replicate 3
KS2 C266I
NaS2 C266I

Figure S5

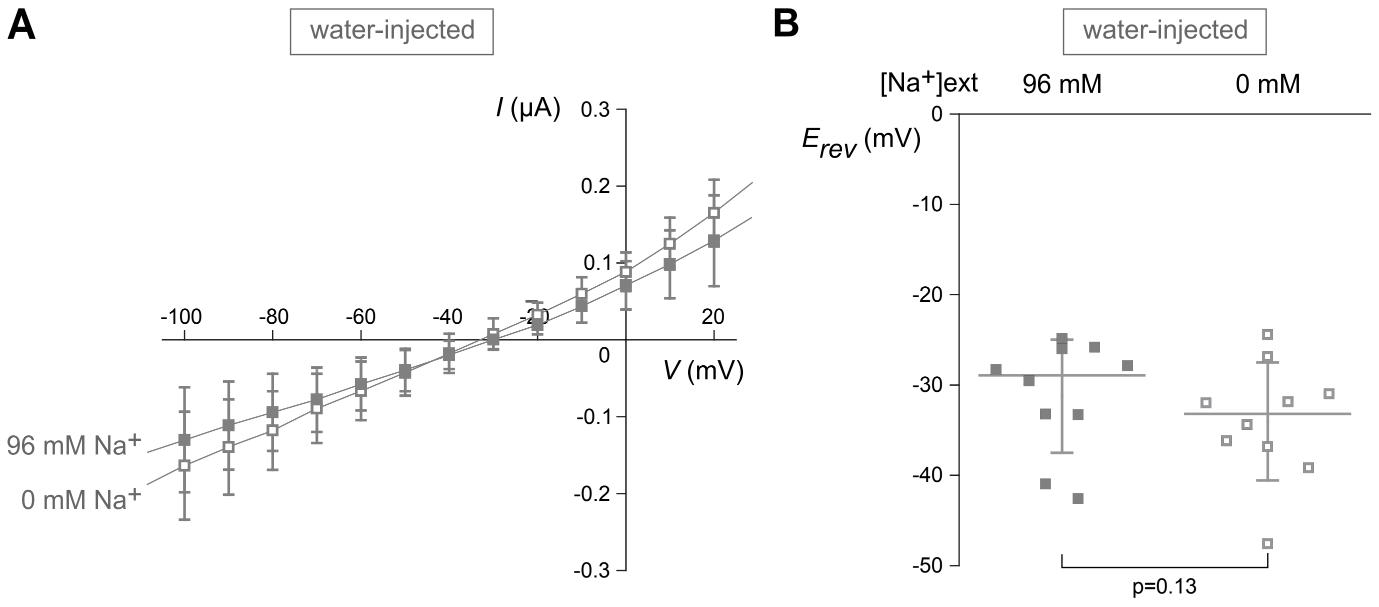
A
water-injected
I (µA)
0.3
0.2
0.1
-0.1
-0.2
-0.3
-100
-80
-60
-40
-20
0
20
V (mV)
96 mM Na+
0 mM Na+
B
water-injected
[Na+]ext
96 mM
0 mM
Erev (mV)
0
-10
-20
-30
-40
-50
p=0.13
